# Degradation of IKAROS prevents epigenetic progression of T cell exhaustion in a novel antigen-specific assay

**DOI:** 10.1101/2024.02.22.581548

**Authors:** Tristan Tay, Gayathri Bommakanti, Elizabeth Jaensch, Aparna Gorthi, Iswarya Karapa Reddy, Yan Hu, Ruochi Zhang, Aatman Doshi, Sin Lih Tan, Verena Brucklacher-Waldert, Laura Prickett, James Kurasawa, Michael Glen Overstreet, Steven Criscione, Jason Daniel Buenrostro, Deanna A. Mele

## Abstract

In cancer, chronic antigen stimulation drives effector T cells to exhaustion, limiting the efficacy of T cell therapies. Recent studies have demonstrated that epigenetic rewiring governs the transition of T cells from effector to exhausted states and makes a subset of exhausted T cells non-responsive to PD1 checkpoint blockade. Here, we describe an antigen-specific assay for T cell exhaustion that generates T cells that are phenotypically and transcriptionally similar to those found in human tumors. We performed a screen of human epigenetic regulators, identifying and validating IKAROS as a driver of T cell exhaustion. We found that the IKAROS degrader iberdomide prevents exhaustion by blocking chromatin remodeling at T cell effector enhancers and preserving binding of AP-1, NF-κB, and NFAT. Thus, our study uncovered a role for IKAROS as a driver of T cell exhaustion through epigenetic modulation, providing a rationale for the potential use of iberdomide in solid tumors to prevent T cell exhaustion.

**Highlights:**

- Novel *in vitro* assay generates antigen-specific exhausted T cells from human T cells
- IKAROS (IKZF1) is a key driver of T cell exhaustion
- Iberdomide (IKZF1/3 degrader) prevents the progression of exhaustion
- TF footprinting reveals IKAROS silences effector genes by inhibiting AP-1, NF-κB and NFAT binding

## Introduction

In cancer patients, tumor specific T cells are driven to exhaustion in the tumor microenvironment and become non-functional. Immune checkpoint therapies directed at blocking inhibitory receptors have shown great promise across several tumor types, albeit effective only in a fraction of patients. Exhaustion of T cells is also a significant problem for chimeric antigen receptor (CAR) T and TCE (T cell engager) therapies^1,2,3,4^. Functional genomics has defined a central role for epigenetics in driving and maintaining exhausted T-cell states. Distinct alterations to DNA methylation^5^ and chromatin accessibility^6^ profiles at well-defined immunological loci accompany T cell exhaustion, which includes stable silencing of effector cytokines necessary for T-cell mediated killing. Single-cell ATAC-seq studies reveal that exhausted T cells are heterogenous, displaying differences in their function^7,8^ and in their response to therapy^8,9^.

Epigenetically reprogramming exhausted T cells to a responsive state could enable broader and perhaps deeper responses to cancer immunotherapies across indications. Towards this end, epigenomic modifications such as pharmacologic inhibition of DNA demethylation^5^, histone deacetylases^10,11^, SWI/SNF^12,13^, and EZH2^14,15,16^ have been reported to rescue aspects of T cell exhaustion. However, current understanding of T cell exhaustion comes mostly from chronic viral infection and syngeneic tumor studies in mice. A model of human T cells exhausted in a physiologically relevant manner would greatly enhance understanding of this process and would enable screening of relevant agents for human cancer immunotherapy.

We sought to identify and expand our knowledge of key drivers in T cell exhaustion using human T cells. We first developed a novel human T cell exhaustion model using healthy donor PBMCs (peripheral blood mononuclear cells) to generate exhausted antigen specific T cells which demonstrated the hallmarks of exhaustion: lower cytotoxicity and cytokine secretion. We characterized these exhausted T cells using single-cell epigenome and transcriptome profiling, demonstrating that they resemble human tumor infiltrating lymphocytes (TILs). We then performed a CRISPR screen in human T cells exhausted by repeated stimulation and identified IKZF1 (IKAROS) as a key player in establishing T cell exhaustion. Further, we demonstrate that T cell exhaustion can be prevented by treating cells with iberdomide, an IKAROS degrader in clinicals trials to treat multiple myeloma^17^. Transcriptional profiling and transcription factor (TF) footprinting analysis revealed that iberdomide halts IKAROS-driven chromatin remodeling at key T cell effector enhancers, preventing T cell exhaustion phenotypes. Our study describes a novel role for IKAROS in T cell exhaustion and provides a mechanistic understanding of how iberdomide prevents exhaustion and maintains a functional T cell state.

## Results

### A novel antigen-specific exhaustion model generates exhausted T cells consistent with TILs from human tumors

To investigate the molecular mechanisms of exhaustion, we developed a novel antigen-driven model of T cell exhaustion that closely mimics the *in vivo* process. We sought to use peptide MHC driven stimulation of T cells to mirror the activation stimulus of T cells in the tumor instead of non-specific anti-CD3/CD28 activation methods which bypass antigen presentation to directly stimulate the T cell receptor (TCR)^18,19^ We utilized known MHC class I peptides from viral antigens (CMV, Influenza and EBV) to expand virus specific T cells from healthy donor PBMCs containing memory T cells from prior exposures (Fig. 1A).

**Figure 1.**
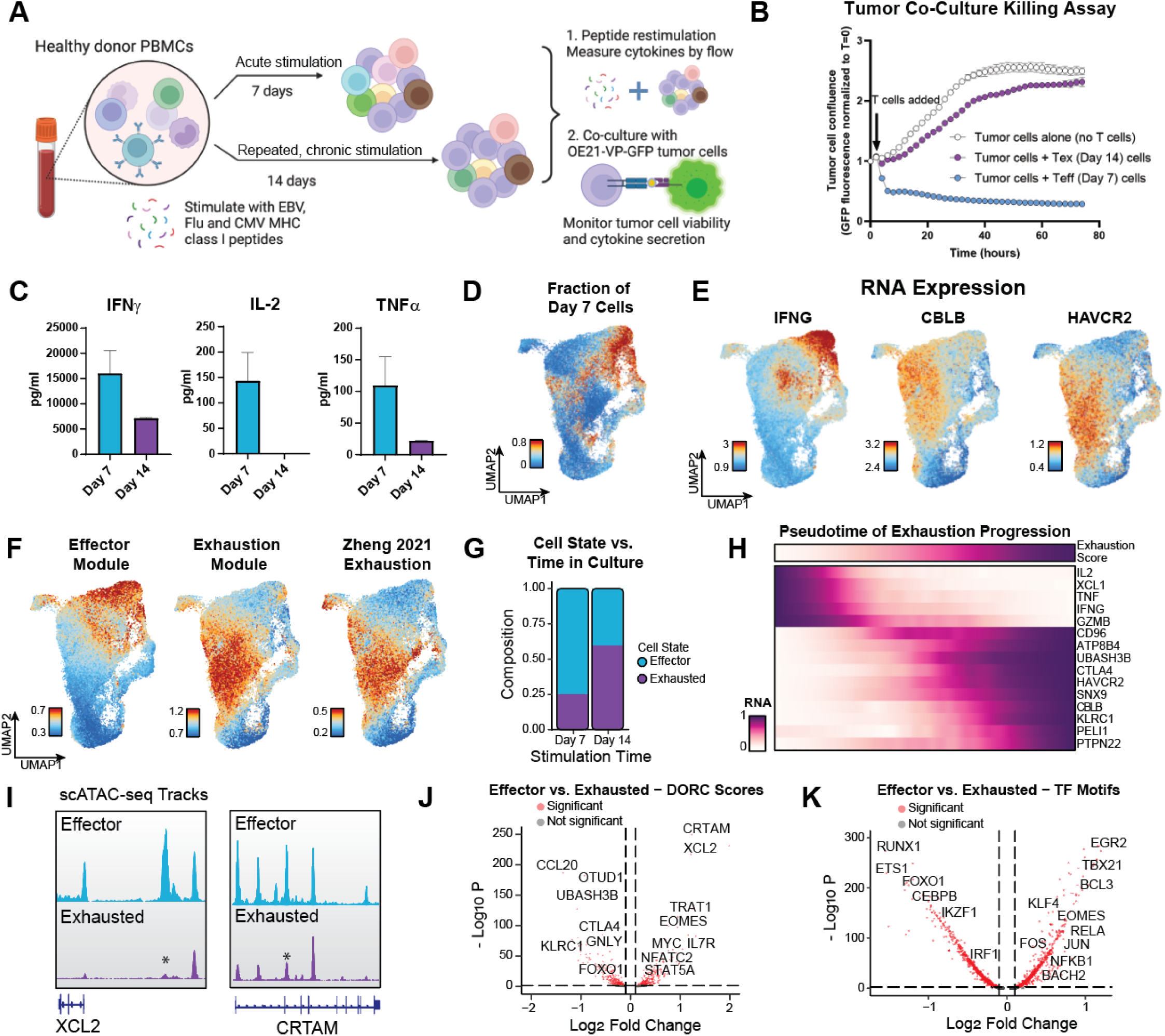
A novel human antigen specific model for T cell exhaustion. A) A schematic of the novel exhaustion assay using peptide stimulation. Figure created using BioRender.com B) Tumor: T cell coculture showing loss of cytotoxic ability of day 14 cells. C) Cytokine analysis for IFNγ (left), IL-2 (center), and TNFα (right) of culture supernatants measured by MSD. D) Distribution of day 7 cells in UMAP generated using RNA expression profiles. E) Gene expression of marker genes on UMAP. F) Gene expression signatures of effector or exhausted T cells called in current dataset or Zheng *et al.* 2021. G) Composition of effector or exhausted T cells by antigen stimulation time. H) Gene expression for individual genes in cells sorted by combined exhausted score. I) ATAC-seq tracks aggregating scATAC-seq profiles from top 5% of most exhausted or most effector cells. Adjusted p-values are 1.2 * 10^-48^ for XCL2 and 1.7 * 10^-23^ for CRTAM. J) Differential accessibility at Domains of Regulatory Chromatin (DORC) associated with specific genes. Significance thresholds are absolute value log2FC > 0.1 and p-value < 0.01. K) Differential TF motif activity calculated using chromVAR scores. Significance thresholds are absolute value log2FC > 0.1 and p-value < 0.01.

Upon peptide re-challenge, T cells under chronic repeated stimulation for 14 days (Tex) had lower levels of cytokine production than cells subjected to acute stimulation for 7 days (Teff), a classic sign of T cell exhaustion (Suppl. Fig. 1A). To assess the cytotoxic ability of day 7 and day 14 cells, we engineered an OE21 tumor cell line that expressed GFP and the same viral peptides used for expansion (Suppl. Fig. 1B). By co-culturing viral peptide expanded T cells with the engineered OE21-viral peptide-GFP (OE21-VP-GFP) tumor cell line, we were able to assess tumor killing in an antigen dependent manner. Day 7 cells were able to efficiently kill tumor cells in contrast to the day 14 cells which were unable to control tumor cell growth, supporting an exhaustion phenotype for day 14 cells (Fig. 1B, Suppl. Fig. 1C). Culture supernatants from the tumor cell co-culture revealed reduced cytokine secretion by day 14 cells (Fig. 1C) consistent with their overall reduced functionality.

To comprehensively characterize our antigen-specific model, we sought to determine if our exhausted T cells recapitulated exhaustion-associated cell signatures seen in previous studies^6–9^. To do this, we enriched antigen-specific T cells from day 7 and day 14 cultures (Suppl. Fig. 1D) with flow sorting using MHC tetramers staining for viral peptide reactive T cells. We performed SHARE-seq, a multimodal single-cell ATAC and RNA sequencing method^20^, and generated 84,685 matched ATAC and RNA profiles of antigen-specific cells across four donors with median ATAC and RNA depths of 11,772 fragments and 2,730 reads per cell, respectively (Suppl. Fig 1E, Suppl. Fig. 1F). After Harmony^21^ correction to remove donor-specific batch effects, uniform manifold and projection (UMAP) visualization of the RNA profiles reveal that T cells separate by length of stimulation time (Fig. 1D). This separation of day 7 cells is mirrored by the higher expression of cytokines such as IFNG, IL2, TNF, and GZMB that are essential for effector function and the absence of expression of known exhaustion regulators such as CBLB^22^, HAVCR2, SNX9^23^, and ENTPD1^24^ (Fig. 1E).

To systematically map the differential gene programs within day 7 (effector) and day 14 (exhausted) cells, we created gene modules based on covarying expression profiles across clusters of cells (Suppl. Fig. 2A). We detected four modules that we annotated as effector, exhaustion, transition, and memory modules (Suppl. Fig. 2B, Suppl. Fig. 2C). The effector module was annotated based on canonical cytokine markers IFNG, IL2, TNF and had top GO annotations that included positive regulation of metabolic process and regulation of leukocyte activation, consistent with the effector phenotype of high proliferation and activation (Suppl. Fig. 2D). Meanwhile, the exhaustion module included negative regulators of T cell activation such as HAVCR2, CBLB, and CTLA4 and had top GO annotations such as negative regulation of T cell proliferation and T cell activation. We labelled the memory module based on marker genes such as IL7R and LEF1 typically associated with naïve or memory T cells. Interestingly, cells expressing this module had no canonical markers of activation or exhaustion despite having a TCR that recognizes viral antigens. Lastly, we labeled the transition gene module because it includes both effector and exhaustion associated genes such as CRTAM, REL, and NR4A3 not included in other signatures. These genes may associate in the same module because they persist longer in transitioning effector cells or turn on early in initiating exhaustion progression.

We compared our de novo gene modules to gene programs found in human cancer CD8+ TILs^25^ to determine the biological relevance of our antigen-specific model (Suppl. Fig. 2E). Our exhaustion gene module was highly enriched in the same genes comprising the terminally exhausted cluster (p = 1.83 * 10^-45^) found in multiple tumor types, supporting that our exhausted T cells reflect their *in vivo* clinical counterpart. Indeed, the expression of our exhaustion module highly overlapped with the expression of the exhaustion program found in the human cancer CD8+ TILs^25^ at the single-cell level (Fig. 1F). On the other hand, our effector gene module had little enrichment with any TILs cluster, which is expected due to the dysfunctional state of T cells in patient tumors. Our memory module had high enrichment with terminally differentiated effector memory T cells (p = 1.15 * 10^-19^), GZMK+ effector memory T cells (p = 1.61 * 10^-18^), and ZNF683+ CXCR6+ resident memory T cells (p = 5.25 * 10^-25^). Therefore, our model produces an antigen-specific memory T cell population and a spectrum of exhausted cell states recapitulating the functional and transcriptomic features found in exhausted T cells from human tumors.

To analyze the progression and heterogeneity of exhaustion in our model, we took advantage of the single-cell resolution of our dataset and compiled a score reflecting the overall T cell state of exhaustion (see methods). Low-scoring effector cells were significantly enriched in day 7 T cells while high-scoring exhausted cells were enriched in day 14 T cells (P-value < 2.2e-10, Kolmogorov Smirnov test) (Fig 1G). Despite this enrichment, however, almost 45% of day 14 T cells still retained an effector-like transcriptomic signature, highlighting the heterogeneity of exhaustion and the need for single-cell measurements. We used this score to rank cells and visualize the trajectory from effector to exhaustion in our assay (Fig 1H). Interestingly, genes like KLRC1 are only expressed in terminally exhausted cells while other exhaustion genes like CBLB are expressed throughout the transition of effector cells to exhaustion. This reflects the role of CBLB as a key inhibitor of TCR signaling and early regulator of T cell exhaustion^26^. These results are consistent with the acquisition of an exhaustion-associated transcriptional signature over the course of our assay.

We also investigated chromatin profiles in the day 7 and day 14 cells to identify peaks uniquely accessible in effector or exhausted cells. Key enhancers of cytokine genes at XCL2 and CRTAM have decreased chromatin accessibility in exhausted cells (Fig. 1I). Interestingly, enhancers at IFNG (p = 0.0178) and IL2 (p = 0.0424) were not significantly silenced in day 14 cells, indicating that loss of expression during exhaustion progression may be attributable to other factors such as differential transcription factor binding (Suppl. Fig. 2F). To broadly quantify gene-associated changes of chromatin accessibility, we correlated the paired RNA expression of each gene with the chromatin accessibility of all peaks to identify gene-enhancer relationships. Genes with high numbers of regulating enhancers are defined as Domains of Regulatory Chromatin (DORCs)^20^ and are enriched for key exhaustion regulators such as NR4A3 and CTLA4 (Suppl. Fig. 2G). We calculated DORC scores by summing the accessibility of regulating enhancers for each gene. DORC scores were decreased in exhausted cells for genes implicated in maintaining T cell effector function such as CRTAM (p = 3.25 * 10^-249^), EOMES (p = 3.38 * 10^-99^), and NFATC2 (p = 4.11 * 10^-35^), in addition to IFNG (p = 2.85 * 10^-11^) and IL2 (p = 3.92 * 10^-5^) as cells progress along the exhaustion trajectory (Fig. 1J). On the other hand, DORC scores were increased for exhaustion-associated genes such as at CTLA4 (p = 1.64 * 10^-84^), UBASH3B^33^ (p = 4.90* 10^-152^), and KLRC1^34^ (p = 1.58* 10^-68^).

To identify transcription factor regulators of exhaustion, we computed TF motif accessibility scores^27^. KLF4, NF-κB1/REL, and JUN/FOS transcription factor families had higher activity in effector T cells, consistent with previous data describing these TFs as important drivers of T cell activation^28,29^ (Fig 1K). On the other hand, exhausted T cells had higher activity in the transcription factors FOXO1^30^ and IKZF1^31^ previously shown to negatively regulate cytokine production. Together, this data demonstrates that this *in vitro* antigen-specific exhaustion assay generates T cells with functional and cell-intrinsic characteristics of exhausted T cells from human tumors and can nominate transcription factors driving exhaustion in an unbiased manner.

### IKAROS (IKZF1) identified as a top hit in a human Tex cell CRISPR KO screen

We set out to perform a CRISPR screen of human epigenetic regulators to find new drivers of T cell exhaustion. However, the antigen-specific assay generates a limited number of exhausted T cells making it technically unfeasible to have appropriate representation of library members. To overcome this limitation, we used a chronic stimulation model using repeated stimulation with anti-CD3/CD28 antibodies more suited for high-throughput screening (Fig. 2A). In this model, T cells produced lower levels of cytokines consistent with exhaustion over successive rounds of stimulation (Fig. 2B, Suppl. Fig. 3A). We included biological replicates and found that across the 3 separate donors, the lethal controls resulted in cell death (Suppl. Fig. 3B) and knockout of beta-2-microglobulin was detectable by flow cytometry (Suppl. Fig. 3C), supporting a functional CRISPR protocol.

**Figure 2.**
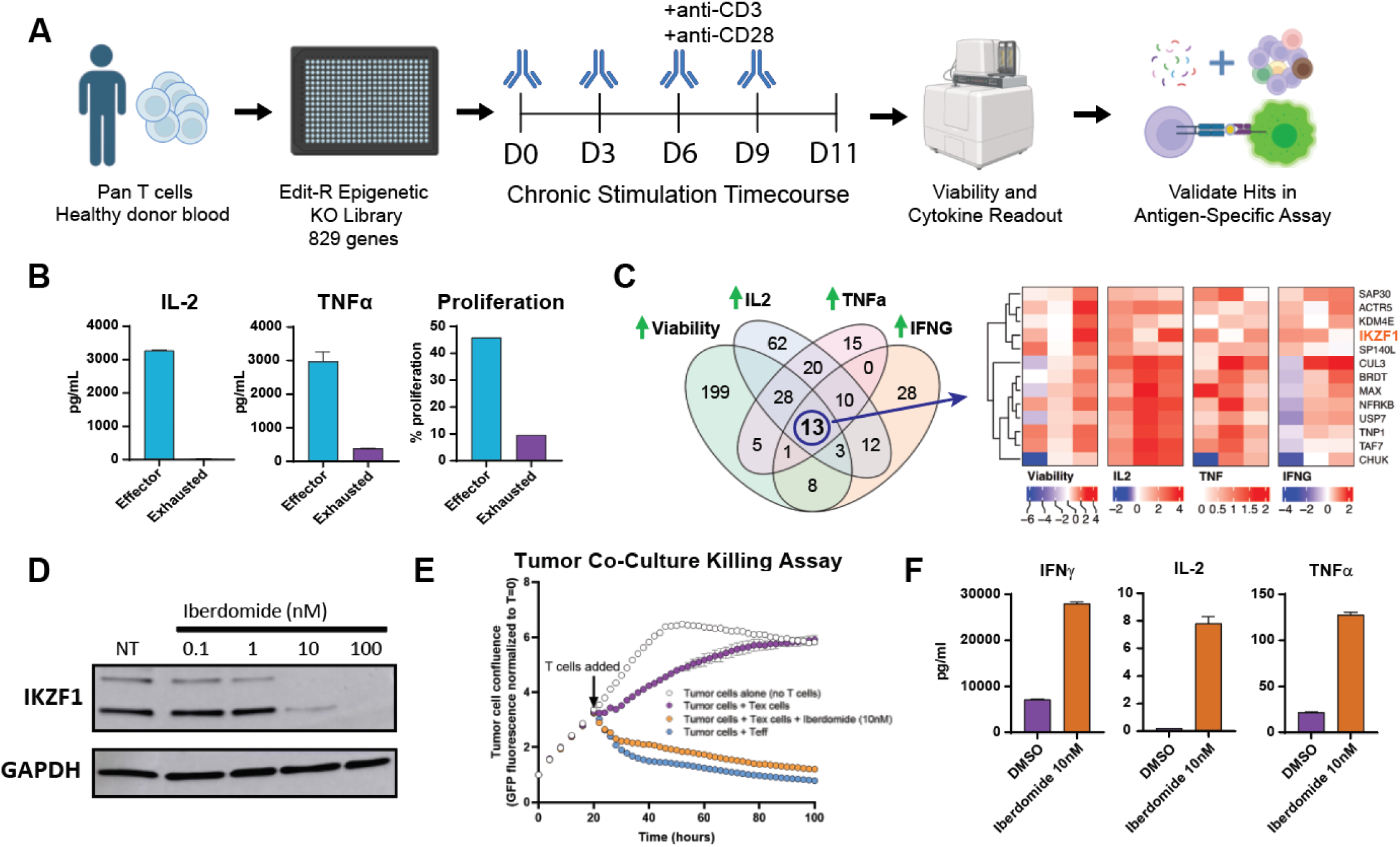
A CRISPR screen in exhausted T cells identifies IKAROS as driver of T cell exhaustion. Iberdomide, a clinical IKAROS degrader, prevents T cell exhaustion. A) Schematic of the CRISPR screen workflow using Immunocult reagent to generate exhausted T cells. B) T cells lose their ability to proliferate and produce cytokines upon repeated chronic stimulation. Cytokines were measured by Meso Scale Discovery (MSD) analysis of cell culture supernatants. Proliferation was measured by detection of EdU incorporated into cells by flow cytometry. C) Top hits from the CRISPR screen. Heatmap of 13 hits with increased cytokine expression and viability shown. D) Western blot showing IKAROS degradation after 7 days of iberdomide treatment. E) Iberdomide treatment of cells during chronic stimulation prevents them from losing E) cytotoxic ability and F) cytokine secretion activity. Cytokine analysis of culture supernatants measured by MSD.

This screen revealed known and novel regulators of T-cell exhaustion. Knockouts of DNMT3a and HDAC2, both of which have previously been reported to have a role in T cell exhaustion^5,10,11,32^, enhanced IL-2 production in Tex cells (Suppl. Fig. 3D), giving us confidence in the success of our screen. Top screen hits also included CUL3^33^, which is a known regulator of T cell function, and ACTR5 and NFRKB, members of the INO80 complex which has been implicated in other exhaustion screens^13^. Other hits such as USP7, a ubiquitinyl hydrolase whose other family members are negative regulators of T cell activation^34^, and CHUK (IKKα), a known inhibitor of inflammation and the NF-κB complex in macrophages^35,36^, were previously unlinked to T cell exhaustion. Finally, IKAROS (IKZF1) was nominated as an exhaustion regulator (Fig. 2C), a transcription factor previously known for its role in lymphocyte differentiation and repressive effect on T cell activation^31,37,38^. We had high confidence in IKAROS as an exhaustion regulator because of its higher activity in exhausted cells (see Fig. 1K) and its known recruitment of histone deacetylase complexes, including the components HDAC2 and SAP30 that were also hits in our screen. Interestingly, cereblon E3 ligase modulators (CELMoDs), currently in trials for multiple myeloma^17^ and acute myeloid leukemia^39^, are known to activate T cell function by inducing the degradation of IKAROS and its family member Aiolos (IKZF3) through cereblon E3 ligase^40^. Additionally, the previous-generation IKAROS degrader lenalidomide has shown beneficial effects in enhancing CAR-T cell function in a small clinical trial^41^. Indeed, upon follow-up CRISPR knockout of IKAROS, the loss of IKAROS protein (Suppl. Fig. 3E) was accompanied by a corresponding increase in cytokine secretion (Suppl. Fig. 3F) even after chronic stimulation, confirming a role for IKAROS in promoting T cell exhaustion and warranting future mechanistic investigation using our assay.

To extend our findings in IKAROS to clinically relevant therapies, we tested if the CELMoD iberdomide (CC-220)^42^ could prevent T cell exhaustion in our physiologically relevant antigen-specific assay. Iberdomide degraded IKAROS within a few hours of treatment in day 7 T cells and maintained negligible IKAROS levels for the remainder of the 14-day chronic stimulation time course (Fig. 2D). Treatment with iberdomide prevented T cell exhaustion phenotypes seen after 14 days of chronic stimulation, restoring tumor cell killing in our coculture assay (Fig. 2E, Suppl. Fig 3G) and increasing cytokine production (Fig. 2F, Suppl. Fig 3H). These results confirmed that iberdomide treatment phenocopies IKAROS knockout and highlight iberdomide as a promising therapeutic to rescue T cell exhaustion.

### Iberdomide preserves the chromatin landscape of effector cells despite chronic stimulation

Although iberdomide was able to sustain effector T cell function in chronically stimulated T cells, we wanted to investigate whether these protective effects extended to the genomic level. We performed SHARE-seq^20^, a paired single-cell RNA and chromatin accessibility sequencing method, on antigen-specific day 14 T cells at the end of the iberdomide and chronic stimulation time course. The iberdomide-treated cells clustered separately from the DMSO-treated cells, reflecting the dramatic effect of the drug (Fig. 3A). In agreement with functional assays, iberdomide treatment upregulated expression of effector genes such as IFNG, IL2, and TNF while downregulating exhaustion markers including TIGIT, CTLA4, and CBLB (Fig. 3B). Interestingly, iberdomide also induced higher IKZF1 expression, likely caused by a compensatory mechanism to IKAROS degradation in cells. We ranked day 14 DMSO or iberdomide-treated cells by their exhaustion score and discovered significant enrichment of iberdomide-treated cells in the effector population (P-value < 2.2e-10, Kolmogorov Smirnov test) (Fig 3C). Treatment with iberdomide also altered the expression of non-exhaustion genes, including programs associated with cellular response to stress (p = 6.88 10^-10^) and defense responses (p = 2.73 * 10^-9^) (Suppl. Fig. 4A).

**Figure 3.**
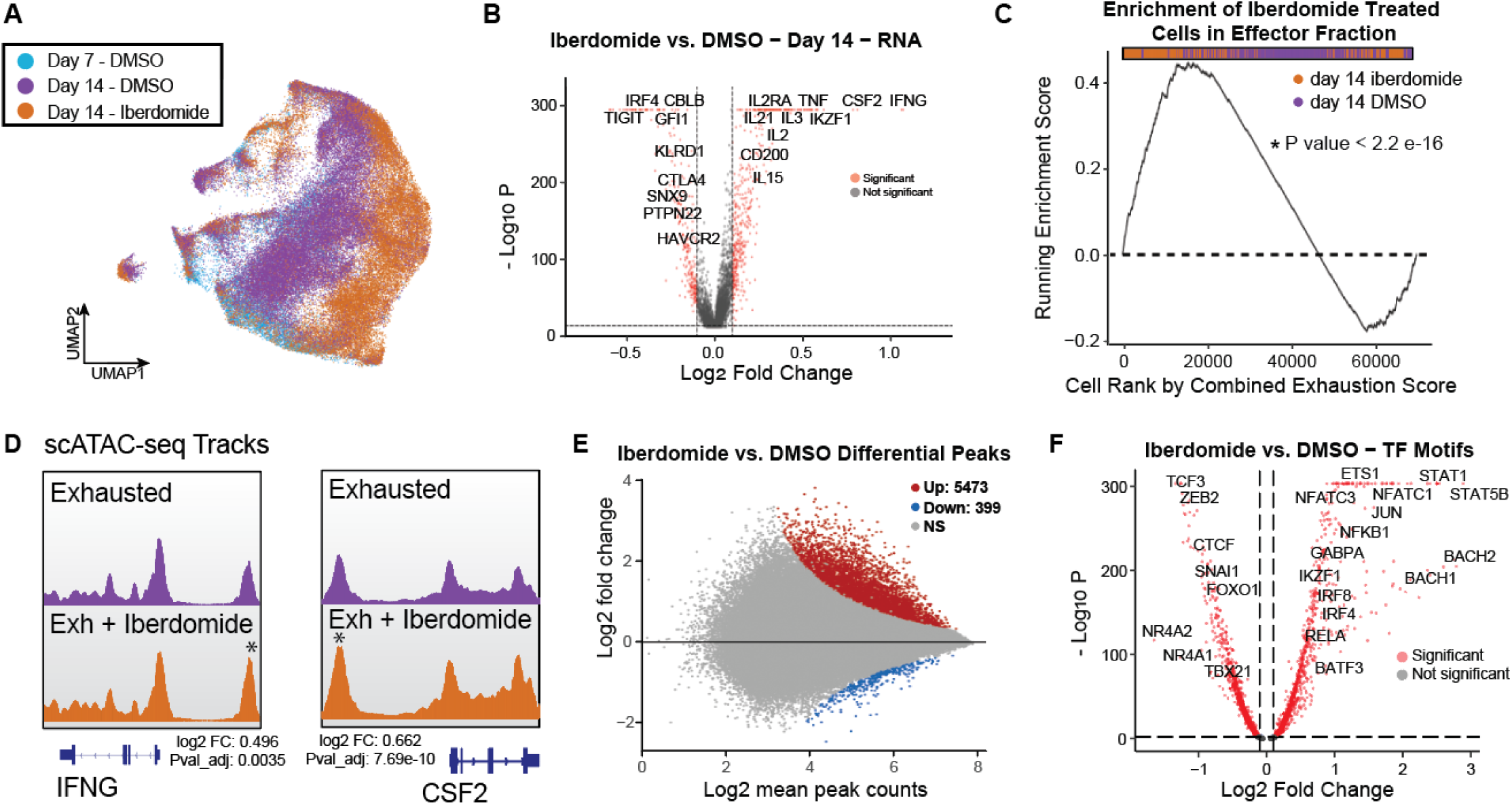
Iberdomide affects gene expression of key effector/regulator genes in Tex cells. A) Donor corrected UMAP of RNA expression of iberdomide and DMSO treated cells. B) Differential RNA expression in T cells treated for 14 days with iberdomide versus DMSO. C) Cumulative enrichment analysis of iberdomide-treated cells with higher effector function. D) ATAC-seq tracks aggregating scATAC-seq profiles from exhausted cells treated with DMSO or iberdomide. E) MA plot of peaks with differential chromatin accessibility in scATAC-seq profiles after iberdomide treatment. F) Differential TF motif accessibility in iberdomide vs. DMSO treated exhausted cells (chromVAR).

Further investigation of chromatin profiles revealed that iberdomide preserves chromatin accessibility at enhancers of immune genes such as TNFSF8 and STAT4 (Fig 3D), but did not affect chromatin accessibility at other key effector genes such as IFNG and IL2. Interestingly, the majority of chromatin changes following iberdomide treatment were gains in accessibility, suggesting that IKZF1 is a repressor of chromatin accessibility (Fig 3E). To identify how iberdomide-induced IKAROS degradation restores T-cell exhaustion, we used chromVAR to globally detect TF motifs with differential accessibility. Motifs of TF families associated with effector function such as JUN/FOS, IRF, REL/ NF-κB, and NFAT were significantly more accessible after iberdomide treatment (Fig 3F). On the other hand, the transcription factors NR4A1 and NR4A2, known to promote T cell exhaustion^43^, had reduced activity after treatment, indicating that IKAROS function was necessary for the onset of the exhaustion program.

To further associate changes in transcription factor function to target genes, we analyzed iberdomide-induced differentially expressed genes using the Causal Reasoning algorithm^44^. Briefly, Causal Reasoning uses protein-protein interaction databases to infer direct relationships from gene expression data, identifying the top genes mediating iberdomide response. Consistent with our chromatin accessibility analysis, the transcription factors NFATC2, REL, and IRF8 scored among the top factors inducing changes in expression of key effector cytokines (Suppl. Fig. 4B). Activity of these TF families is associated with T cell effector function in both viral and tumor backgrounds^45^, suggesting that IKZF1 may function to repress these transcriptional programs. Together, analysis of the transcriptome and epigenome provides further mechanistic understanding of iberdomide’s ability to preserve T cell effector function, despite chronic stimulation.

### Iberdomide Rescues Transcription Factor Binding at Effector Enhancers

To identify the primary mechanism of iberdomide in preserving T cell effector state, we acutely treated primary human T cells with iberdomide for 6 hours, stimulated them using Immunocult anti-CD3/CD28 reagent, and performed bulk ATAC-seq^46^. Of the regions with differential chromatin accessibility, 95.1% had higher accessibility, affirming that IKAROS functions as a repressor of its target genes (Fig. 4A). This differential ATAC-seq peak set had the highest similarity with IKAROS ChIP-seq sites when compared to Cistrome DB’s database of all ChIP-seq studies^47^, confirming that acute depletion reveals the direct effects of IKAROS degradation (Suppl. Fig 4C).

**Figure 4.**
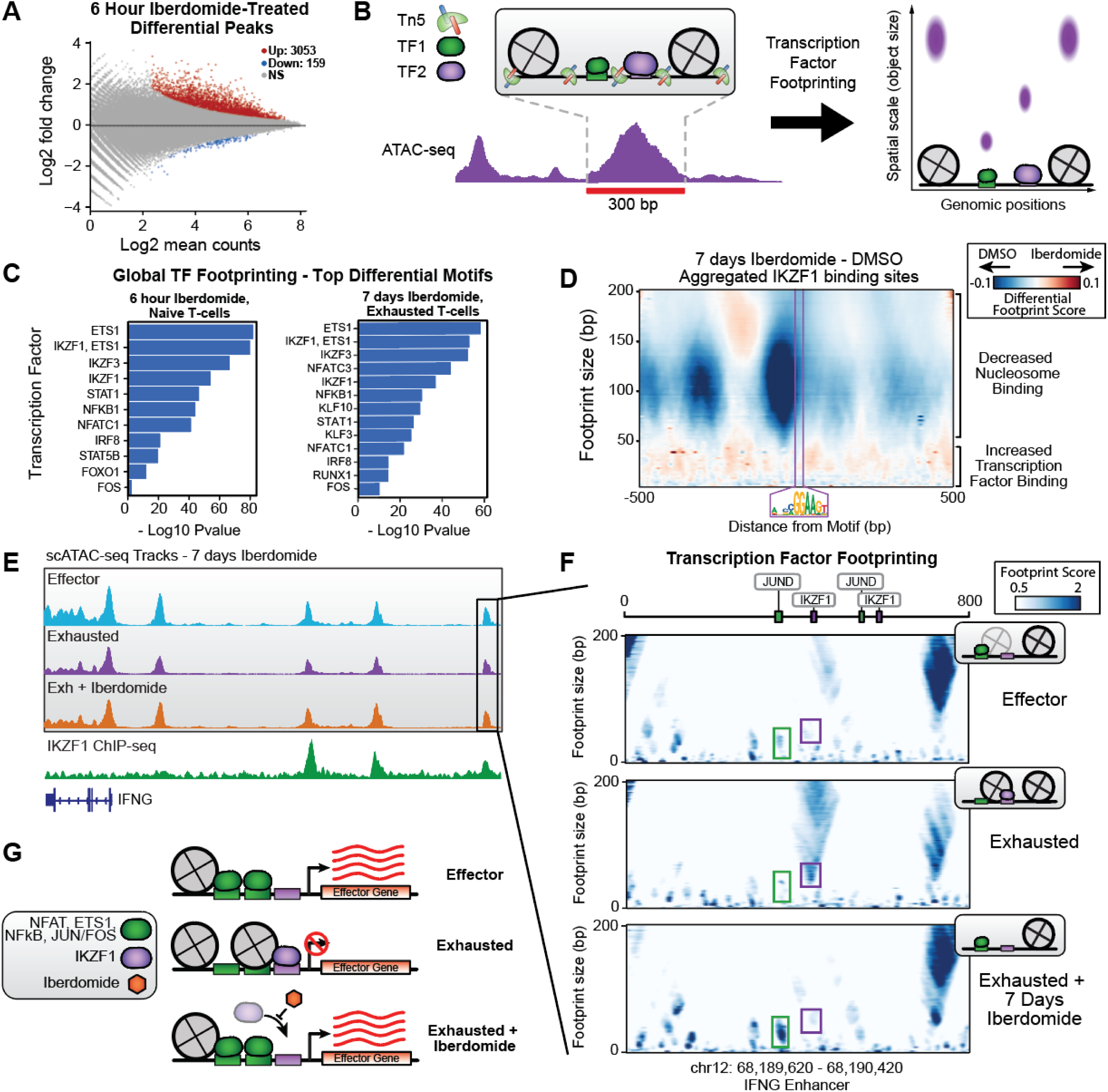
TF footprinting analysis reveals loss of nucleosomes and increased AP-1, NF-κB and NFAT Binding at effector enhancers following iberdomide treatment. A) MA plot of peaks with differential chromatin accessibility in bulk ATAC profiles after 6 hour iberdomide treatment. B) Schematic of transcription factor footprinting analysis. C) Top motifs with differential binding scores across all peaks in bulk ATAC (left) or scATAC-seq (right). D) Aggregated multiscale footprinting plots of peaks containing IKZF1/ETS1 motif and filtered for the highest 25% in IKZF1 Chip-seq signal. E) ATAC-seq tracks aggregating scATAC-seq profiles from untreated effector, untreated exhausted cells, or exhausted cells treated with iberdomide. IKZF1 ChIP-seq tracks from Sun et. al 2022 in purple (bottom). F) Multiscale TF footprinting in enhancer marked in (E) from effector cells, exhausted cells, or iberdomide-treated exhausted cells. Schematics of TF and nucleosome binding inferred from plots shown below each plot. G) Model of iberdomide mechanism of action in restoring effector function.

To further investigate the interplay of transcription factors at exhaustion-associated regulatory elements, we employed TF footprinting^48^. By looking at individual Tn5 cuts rather than smoothing over peaks, TF footprinting allows resolution of sub-enhancer events including individual DNA element binding events. We use TF footprinting to identify differential nucleosome or TF binding events even in enhancers with no change in chromatin accessibility, giving us greater resolution into the mechanism of iberdomide-mediated maintenance of T cell effector function (Fig 4B). In both the acute 6 hour iberdomide treatment on naïve T cells and 7 day treatment on exhausted T cells, TF footprinting revealed that iberdomide increased binding at key motifs of ETS1, JUN/FOS, STAT/IRF, REL/ NF-κB, and NFAT families as was previously seen with chromVAR analysis (Fig 4C). Previous generation immunomodulatory drugs (IMIDs) closely related to iberdomide have been shown to upregulate JUN/FOS and NFAT families^49^, further supporting our results. Differential binding at these key motifs during iberdomide treatment largely occurs in the absence of differential TF expression (see Fig. 3B and Sup. Table 1), highlighting that these differential binding patterns result from IKZF1 degradation reshaping the regulatory element landscape. Additionally, the similarity in affected motifs from Ikaros degradation in naïve vs. exhausted T cells indicate that iberdomide primarily prevents exhaustion by preserving activity of TFs at effector genes, rather than reversing activation of exhaustion genes which would not be relevant in the naïve T cells.

Intriguingly, loss of nucleosome binding upon iberdomide treatment primarily occurs at a GGAA motif shared by IKZF1 and the ETS family of TFs (Suppl. Fig. 4D). IKZF1 has been previously identified as a direct competitor with Ets activators at joint binding sites, giving us confidence in the accuracy of our footprinting analysis^50,51^. The ETS family is implicated with pioneer factor behavior in effector T cells, where it is required for the binding of adjacent activating TFs like AP-1, NFAT, and REL^45^. At peaks with IKAROS ChIP-seq signal^52^, aggregated TF footprinting results illustrated that iberdomide causes a loss of binding of nucleosomes at the joint IKAROS/ETS motif, coupled with a general increase in footprints of transcription factor size (Fig. 4D) when compared to a TSS background control (Suppl. Fig. 4E), in agreement with our previous analysis.

There were specific examples of this nucleosome ejection and increased transcription factor binding at enhancers of key immune loci like IFNG (Fig. 4E). At an enhancer 31 kb from the IFNG promoter with high IKAROS ChIP-seq signal, we previously did not detect any changes in chromatin accessibility. However, we used our new TF footprinting tools to find increased nucleosome binding during exhaustion progression at an IKZF1/3 motif in this enhancer, coupled with loss of Jun binding at a nearby AP-1 motif (Fig 4F). Iberdomide treatment is sufficient to prevent exhaustion reprogramming at this specific enhancer, preserving AP-1 binding and IFNG expression in chronically stimulated T cells. These regions were not nominated by differential chromatin accessibility levels despite the clear ejection of a nucleosome (Suppl. Fig. 4F), highlighting the need for TF footprinting to identify more nuanced changes to sub-enhancer events. At other locations, such as at a distal enhancer of CSF2 (GM-CSF) with differential chromatin accessibility (Suppl. Fig. 4G), there was a similar loss of nucleosome binding upon iberdomide treatment coupled with a gain of transcription factor binding at the joint IKZF1/ETS motif (Suppl. Fig. 4G). At another distal enhancer of IFNG, loss of nucleosome binding at an IKZF1 motif was paired with loss of binding of the transcription factor NR4A2 (Suppl. Fig 4H).

This leads us to a model whereby ETS1 and IKZF1 competitively bind during exhaustion progression (Fig. 4G). During exhaustion, increased IKZF1 activity promotes nucleosome occupancy and shutdown of effector gene enhancers containing AP-1, NFAT and REL binding sites. Iberdomide-mediated degradation of IKZF1 preserves ETS binding and effector gene function. However, iberdomide treatment did not affect inhibitory receptor expression (LAG3, PD-1, and TIGIT) as measured by flow cytometry analysis (not shown). In summary, we propose that the direct impact of iberdomide in preventing exhaustion progression is achieved by removing the inhibitory effects of IKZF1 on effector gene function.

## Discussion

Epigenetics controls the differentiation of T cells into specialized subsets, including exhausted T cell states which have been associated with the heterogeneity of patient response rates to cancer immunotherapy^8,9^. However, the field lacks a comprehensive view of epigenetic remodeling during exhaustion progression and whether therapies can be developed to revert the epigenetic barriers establishing an exhaustion cell state.

Here, we identified a novel role for IKAROS as a driver of T cell exhaustion, in addition to its previously known roles in T cell development^53^, differentiation^38^, and cytokine suppression^31,54,55^. Using our antigen-specific *in vitro* assay and single-cell multi-omics, we characterized the mechanism of the IKAROS degrader iberdomide across multiple layers of gene regulation, providing an integrated understanding of how IKAROS degradation prevents silencing of effector genes and preserves effector function. We discovered that iberdomide prevents IKAROS-mediated recruitment of nucleosomes, safeguarding chromatin accessibility and binding of the TF families NFAT, AP-1, ETS, and NF-kB/REL at effector genes despite chronic stimulation. Additional experiments are required to determine the plasticity of these iberdomide-induced changes, especially in the case of treatment withdrawal. Our findings are consistent with previous reports of IKAROS acting as a transcriptional repressor through the recruitment of histone deacetylase (HDAC) complexes^56,57^. Indeed, our CRISPR screen identified HDAC components (SAP30 and HDAC2) as regulators, raising the possibility that HDAC inhibitors may phenocopy or amplify the effects of iberdomide treatment.

Although iberdomide is a more specific degrader than previous generation IMIDs, it degrades both IKAROS (IKZF1) and a closely related family member AIOLOS (IKZF3). In our ATAC-seq dataset from iberdomide treated T cells, our list of differential peaks had the highest overlap with IKAROS ChIP-seq datasets, suggesting that IKAROS degradation contributes significantly to the observed phenotype. Interestingly, the IMID lenalidomide has been reported to partially rescue T cell exhaustion through AIOLOS degradation^58^. AIOLOS degradation also enhanced CAR-T cell proliferation and activity^59^, suggesting that IKAROS and AIOLOS may have similar roles in T cell exhaustion. Degradation of these two proteins by CELMoDs such as iberdomide impacts the function of multiple cell types beyond T cells^60,61^, however, indicating dose optimization, scheduling, and combination drugs for iberdomide (or other IKAROS degraders) might be needed to translate these *in vitro* findings to clinical benefits in patients.

Despite the necessity of epigenetic remodeling in the progression of exhaustion, targeting epigenetic factors to restore exhausted T cell function has been challenging. Employing pleotropic agents such as Dnmt1i^62^, HDACi^63^ and even more focused inhibitors targeting readers (BETi^)64,65^ and HMT (EZH2i)^14,15,16^ have had limited clinical success in treating tumors. Identification of the TF families Tox and NR4A as exhaustion regulators^66–69^ has been similarly challenging to translate to therapies due to difficulty in druggability. Our identification of a clinical stage drug, iberdomide, and its mechanism in preventing T cell exhaustion, provides a rationale for using iberdomide and other selective IKAROS degraders in the clinic to revive exhausted T cells in solid tumors. Iberdomide may also be promising for preserving effector function of TILs and engineered T cells (CAR-T, TCR-T) during ex vivo expansion when they are subject to long periods of stimulation, highlighting the broad potential use of Ikaros degraders for oncology treatment.

## Supplementary Figures

**Suppl. Fig. 1.**
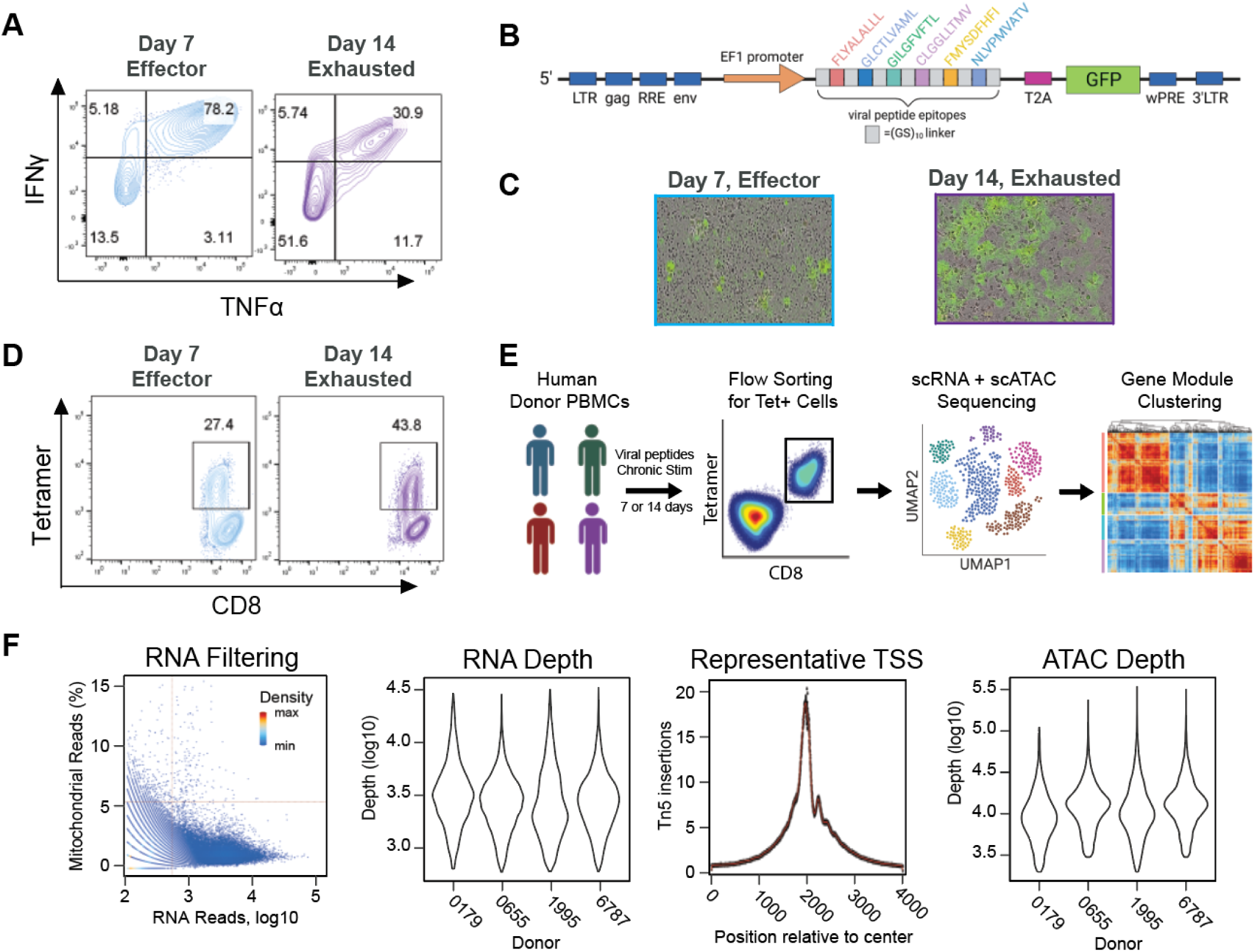
Tetramer Sorting of Antigen-specific T Cells and Data Quality of SHARE-seq Dataset. A) Cells were stained for intracellular cytokines following peptide restimulation of day 7 and day 14 cells. Day 14 exhausted cells show reduced cytokine production. B) Schematic diagram of the construct used in generating the engineered viral peptide and GFP expressing OE21 cell line. C) Representative micrographs showing reduced tumor cell confluence (reduced GFP fluorescence) upon effective tumor cell killing by day 7 effector T cells. D) Flow cytometry plots showing frequency of viral antigen specific (tetramer positive) Teff and Tex cells 7 days and 14 days of culture respectively from an example donor. Depicted events are gated on live CD8 T cells. E) Schematic depicting the extraction of tetramer+ CD8 T cells from human donor blood for multimodal single-cell sequencing. F) QC metrics of SHARE-seq data. scRNA-seq profiles were filtered for < 5% mitochondrial reads and > 500 unique genes (left). Median of 2,730 reads in each profile after filtering (left middle). The enrichment of ATAC reads at transcription start sites (TSS) throughout the genome in scATAC-seq profiles (right middle). Median of 11,772 fragments in each scATAC-seq profile after filtering for > 2000 unique fragments and > 50% fraction of reads in peaks (right).

**Suppl. Fig 2.**
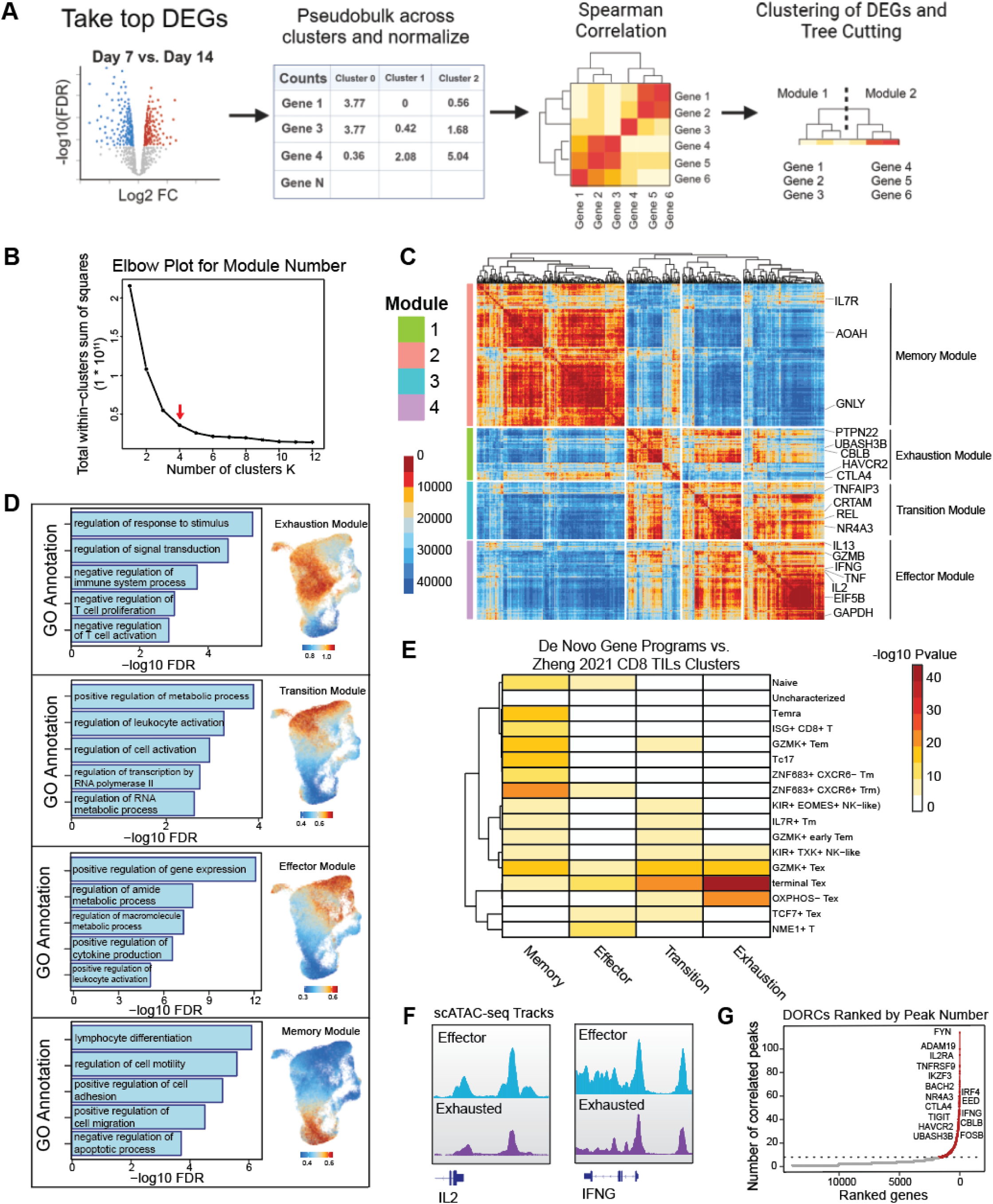
De Novo Gene Programs Capture Full Complexity in Dataset. A) Schematic depicting the creation of de novo gene modules using scRNA-seq profiles. B) Elbow plot determining that four modules minimize the within-clusters sum of squares. C) Heatmap displaying the Euclidean distance between gene expression profiles. The heatmap is broken into four modules based on the tree cutting method (left) and selected genes representing each module are displayed (right). D) Smoothed module scores in UMAP space with accompanying GO annotations. E) Heatmap of pvalues from hypergeometric test comparing overlapping marker genes in our de novo gene modules to Zheng et al. 2021 CD8+ T cell clusters. F) ATAC-seq tracks aggregating scATAC-seq profiles from top 5% of most exhausted or most effector cells for IL2 and IFNG. G) Ranked DORCs by number of correlated peaks. Filtered for at least 8 correlated peaks.

**Suppl. Fig. 3.**
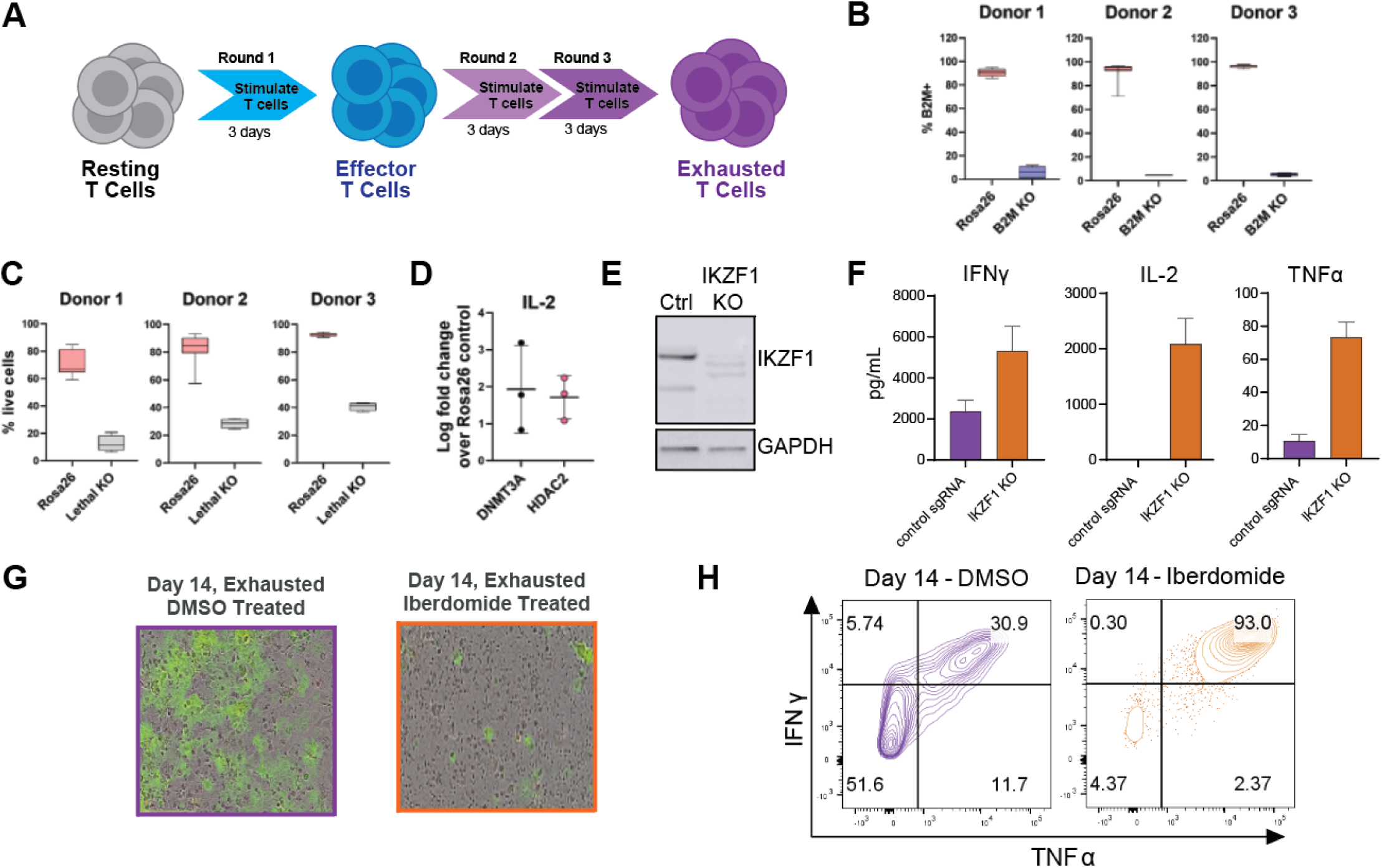
CRISPR Screen Controls and Validation. A) Schematic of repeat stimulation process to generate exhausted T cells. B) Flow cytometry confirmed loss of β2M expression on T cells from three donors upon CRISPR knockout. Each panel shows data from a separate donor. C) CRISPR knockout of essential gene in T cells results in a loss of viability as measured by flow cytometry in cells from three donors. D) CRISPR KO of DNMT3A and HDAC2 resulted in rescue of IL2 secretion across T cells from 3 donors. E) Western blot depicting the degradation of IKZF1 upon sgRNA knockout. F) IKZF1 KO T cells maintain high functional ability upon chronic stimulation. Cytokines measured by MSD from culture supernatants. G) Representative fluorescence micrographs showing enhanced tumor cell killing (reduced GFP fluorescence) when cells are treated with iberdomide during tumor co-culture killing assay. H) Iberdomide treatment of cells during chronic stimulation using the antigen-specific assay prevents them from losing cytokine secretion ability.

**Suppl. Fig. 4.**
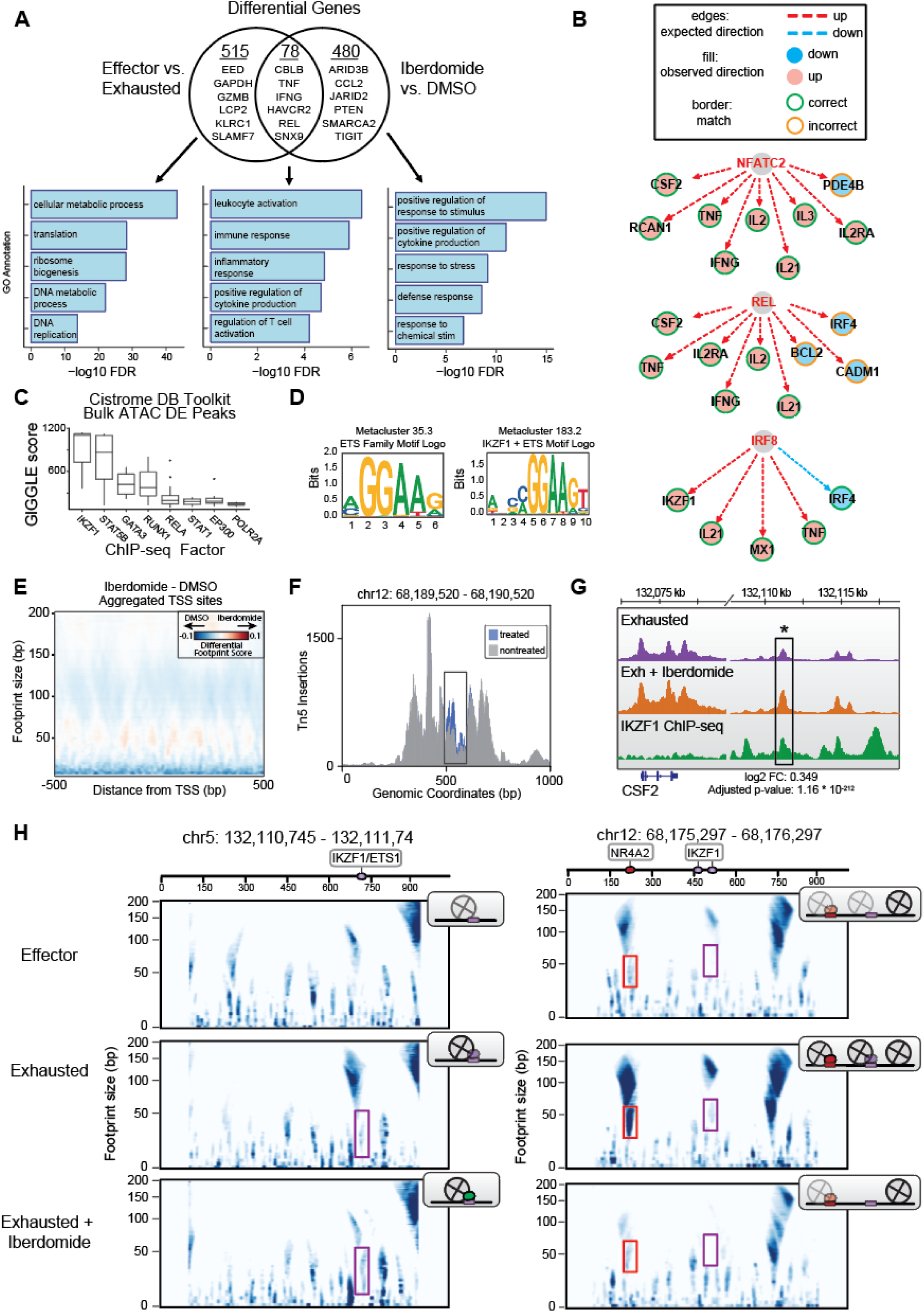
Primary Effects of Iberdomide on Transcription Factor Binding. A) Venn diagram depicting the overlap of differential genes (Pvalue < 10^-100^) comparing Effector vs. Exhausted or Iberdomide vs. DMSO with selected genes displayed. B) Inferred gene regulatory networks for top differential transcription factors in iberdomide vs DMSO. C) Giggle scores calculated for the top 1K differential peaks between 6 hour iberdomide-treated and DMSO-treated cells. Factors were filtered to have at least 3 ChIP-seq datasets to be visualized. D) STAMP consensus motif logos for metaclusters of the ETS family or IKZF1 + ETS. E) Aggregated multiscale footprinting plots of all TSS regions as a background control F) Tn5 insertion plot for iberdomide vs. DMSO treated cells for an enhancer of IFNG shown in Fig. 4E - 4F. G) ATAC-seq tracks for a distal enhancer of CSF2 with differential chromatin accessibility and IKZF1 ChIP-seq signal. H) Multiscale TF footprinting in CSF2 enhancer (left) or IL2 promoter (right) from effector cells, exhausted cells, or iberdomide-treated exhausted cells. Schematics of TF and nucleosome binding inferred from plots shown in top right of each plot. Ikaros binding is replaced by ETS1 binding upon iberdomide treatment.

## Materials and Methods

### Repeat stimulation to induce T cell dysfunction

CD3+ T cells were isolated from healthy donor leukopaks (Stemcell Technologies) using bead separation (Stemcell Technologies) and cryopreserved. Thawed T cells were resuspended in RPMI medium supplemented with heat-inactivated FBS, Pen/Strep, HEPES, non-essential amino acids, sodium pyruvate, Glutamax and 2-mercapto-ethanol (all Gibco). 10 million cells were plated in a T25 tissue culture flask at a density of 1 million/mL. 250 µL of ImmunoCult Human CD3/CD28 T Cell Activator Mix (StemCell Technology) was added to the cells and incubated at 37°C. After 3 days of culture, cells were transferred to fresh culture medium and stimulated again. After 3 rounds of stimulation, cells were washed and stimulated for a final cytokine assessment 24hr post the last round (chronic stim). Freshly thawed cells stimulated for 24hr or 72hr were used for cytokine and proliferation measurements respectively as acute stim control. Cytokines were measured using a Mesoscale Discovery V-PLEX Proinflammatory Panel 1 Human Kit and read on an MSD SECTOR S 600 plate reader. Proliferation of cells in the last 18hr of stimulation was assessed by labelling and staining proliferating cells with Click-iT™ EdU Alexa Fluor™ 647 Flow Cytometry Assay Kit (Thermo Fisher Scientific.) Edu positive cells were detected using a MACSQuant® Analyzer 10 Flow Cytometer (Miltenyi Biotec).

In some experiments, T cells were transferred to P3 buffer (Lonza) and electroporated with NTC (non-targeting control) sgRNA or IKZF1 sgRNA (IDT) and TrueCut Cas9 Protein v2 (Invitrogen). Cells were transferred to culture medium and incubated for 72hr at 37degC. After three rounds of activation with ImmunoCult Human CD3/CD28 T Cell Activator Mix as described above before, cells were washed, counted and 100,000 cells each of control and IKZF1 KO cells plated per well of a 96 well U bottom plate. Cells were stimulated with the activator mix for 24hr and culture supernatants collected for cytokine analysis. KO efficiency was assessed by running cell lysates (72hr post CRISPR editing) on an SDS PAGE gel and western blotting for IKZF1 (clone D10E5, Cell Signalling Technology).

### Arrayed CRISPRko screen in T cells

The screen was performed at Revvity in Cambridge, UK. CD3+ T cells were isolated by magnetic bead separation (StemCell Technology) from PBMCs of three leukopaks (Cambridge Biosciences) and cryopreserved. Revived CD3+ T cells were resuspended in P3 buffer (Lonza) and electroporated (Amaxa HT Nucleofector; Lonza) in a 1:3 v/v ratio of Cas9-NLS (Feldan) and a synthetic sgRNA library targeting 829 genes (3 guides per gene, Revvity Dharmacon). The library used consisted of the Edit-R Human sgRNA Library-Epigenetics (QTE2954349G) and Edit-R Human Synthetic B2M (67), CBLB (8680), Rosa26 (custom synthesis using following gRNA sequence AGTCGCTTCTCGATTATGGG), non-targeting control (NTC) #1, and lethal synthetic sgRNA control #1 (all 3 guides per gene, Revvity Dharmacon). Immediately after electroporation, each well was transferred to RPMI media supplemented with heat-inactivated FBS, Pen/Strep, HEPES, non-essential amino acids, sodium pyruvate, Glutamax and 2-mercapto-ethanol (all Gibco) and splitted into four technical replicates with 20,000 cells per well of a 384 well culture plate. Cells were stimulated with ImmunoCult Human CD3/CD28 T Cell Activator Mix (1:80; StemCell Technology) and incubated at 37°C. Three days after the first stimulation, cells were spun down and supernatant was removed. This was followed by a second, third and fourth round of stimulation every three days with fresh ImmunoCult Human CD3/CD28 T Cell Activator Mix (1:80) resuspended in complete assay medium. Cell Titer Glo 2.0 read-out was assessed on the 2103 EnVision multilabel plate reader (Revvity). Secretion of IL-2, TNF-α and IFN-γ in the supernatant was assessed using HTRF technology (Revvity HTRF) and acquired on the 2105 EnVision multilabel plate reader (Revvity). CBLB served as a positive control that revives exhausted T cell function. Lethal #1, B2M, NTC and Rosa26 served as phenotypical editing controls. For lethal #1, B2M, NTC and Rosa26 cell viability was assessed and surface expression of CD3 and B2M was compared to NTC and Rosa26 using fluorescence-labelled antibodies (anti-CD3 BV421, UCTH1 clone; anti-B2M FITC, 2M2 clone; both BioLegend) and flow cytometry (IntelliCyt iQue Screener Plus; Sartorius) on day 6 post-electroporation. The most significant hits were identified by evaluating genes whose knockout results in a significant (adjusted p value < 0.005) increase in at least 3 out of 4 readouts for at least 2 out of 3 donors.

### Antigen specific exhaustion model

PBMCs were isolated from HLA A2:01 specific healthy donor leukopaks (Stemcell Technologies) using a Stemcell isolation kit and cryopreserved. Cells were thawed and resuspended at 4 million cells/mL in RPMI media supplemented with heat-inactivated FBS, Pen/Strep, HEPES, non-essential amino acids, sodium pyruvate, Glutamax (All Gibco). Cells were plated in a 24 well culture plate (1mL cell suspension/well). 20ng/mL of recombinant human IL-2 (Peprotech) was added to the cells along with 1ug/mL each of HLA A2:01 binding CMV pp65, EBV LMP2, EBV BMLF1, Influenza M peptide and EBV LMP2 viral peptides (MBL International). Cells were incubated for 7 days at 37°C. At the end of 7 days, stimulated cells (consisting of effector T cells) were washed and harvested for further characterization or assays.

To generate exhausted T (Tex) cells, day 7 cells were transferred to fresh culture medium with IL-2 and stimulated again with peptides and 1ug/mL of anti-human CD28 antibody (BD Biosciences) for another 7 days. In some experiments, cells were treated with 10nM iberdomide (MedChemExpress) during this phase of culture. At the end of total 14 days of culture, these chronically stimulated cells (consisting of exhausted T cells) were washed and harvested for further assays/characterization.

Cells were stained with Fixable Live Dead Violet (Thermo Fisher Scientific) followed by staining with fluorescent antibodies for CD3 (clone SK7, Biolegend), CD8 (clone HIT1a, BD Biosciences) and PE conjugated tetramers for HLA-A*02:01 CMV pp65 (NLVPMAVATV), HLA-A*02:01 EBV LMP2 (FLYALALLL), HLA-A*02:01 EBV (GLCTLVAML), HLA-A*02:01 EBV LMP2 (CLGGLLTMV) and HLA-A*02:01 Influenza-M1 (GILGFVFTL) (MBL International). Stained samples were run on a MACSQuant® Analyzer 10 Flow Cytometer (Miltenyi Biotec) to determine the frequency of tetramer positive cells in the sample.

### Peptide restimulation

In some experiments, Day 7 or Day 14 cells (described above) were plated in a 96 well V-bottom plate (200,000 cells/well) and 1ug/mL peptides added to restimulate cells along with Monensin and Brefeldin A (Biolegend). Iberdomide (10nM) was added to some wells as required. Cells were incubated for 4hr, surface stained with fluorescent antibodies for CD3 (clone SK7, BD Biosciences), CD8 (clone SK1, BD Biosciences), CD4 (clone OKT4, Biolegend) followed by intracellular staining for IFNγ (clone B27, BD Biosciences) and TNFα (clone MAb11, BD Biosciences). Samples were run on a BD Biosciences LSRFortessa cell analyzer and data analyzed using Flowjo 10.8 software.

### Generation of viral peptide expressing OE21 cell line

OE21 cells were obtained from the European Collection of Authenticated Cell Cultures (Salisbury, United Kingdom). The sequences for the six viral peptides were concatenated into a single construct, each peptide was separated by a glycine-serine10 linker. This polyepitope insert was synthesized by GeneArt (Waltham, MA), and NheI and NotI restriction sites were used to clone the insert into the pCDH-EF1-MCS-T2A-GFP lentiviral vector (System Biosciences, Palo Alto, CA). OE21 tumor cells were then transduced with the resultant lentivirus, and GFP+ cells were sort-purified.

### Tumor cell killing assay

Viral peptide and GFP expressing OE21 engineered cells from above (OE21-VP-GFP) were seeded at 5,000 cells/well into 96 well plates (PhenoPlate, black, Revvity). Cells were incubated in an Incucyte® S3 Live-Cell Analysis Instrument (Sartorius) overnight. The following day, Day 7 or Day 14 viral peptide expanded PBMCs were added at different E:T ratios after adjusting effector cell number to match tetramer positive cell number in each sample. 10nM iberdomide was added to some wells as required. Plates were returned to the Incucyte and imaged every two hours to monitor tumor cell confluence (GFP fluorescence) over the next 3 days. 24hr after co-culture, 50 µL of culture supernatant was harvested to assess cytokine secretion using a Mesoscale Discovery V-PLEX Proinflammatory Panel 1 Human Kit.

### IKAROS degradation with iberdomide treatment

Human T cells were cultured in medium containing 20ng/mL of IL-2 with different concentrations of iberdomide over 7 days. The extent and kinetics of IKAROS1 degradation was determined by western blotting of cell lysates with an IKZF1 antibody (clone D10E5, Cell Signaling Technology).

### Sorting of antigen specific cells for ShareSeq

To enrich for viral peptide specific CD8 T cells from the Day 7 and Day 14 cells (expanded described above) from 4 separate donors, 10 million expanded cells were plated @ 2 million/mL in a 6-well plate with peptides (1ug/ml) and anti-CD28 (1ug/ml) as described above but without IL-2. 10nM iberdomide was added to some wells for iberdomide treatment cells. After 4 hours of incubation, cells were stained for dead cells (fixable Live Dead stain, Thermofisher), CD8 APC (clone HIT1a, BD Biosciences) and PE conjugated tetramers (MBL International). Cells were fixed by adding formaldehyde (28906, Thermo Fisher Scientific) to a final concentration of 0.2% and incubated at room temperature for 5 minutes and washed in PBS. Cells were kept on ice until sorting. Live tetramer positive cells were bulk sorted into CryoStor freezing medium (07959, STEMCELL Technologies) on a Sony SH800 sorter using a 100µm nozzle and frozen at –80 °C for single cell sequencing.

### SHARE-seq

SHARE-seq was performed as previously described^20^. Briefly, cells were fixed and permeabilized. For joint measurements of single cell chromatin accessibility and expression (scATAC- and scRNA-seq), cells were first transposed by Tn5 transposase to mark regions of open chromatin. To improve transposition, transposition was performed using pre-assembled Tn5 (seqWell, Tagify™ SHARE-seq Reagent). The mRNA was reverse transcribed using a poly(T) primer containing a unique molecular identifier (UMI) and a biotin tag. Permeabilized cells were distributed in a 96-well plate to hybridize well-specific barcoded oligonucleotides to transposed chromatin fragments and poly(T) cDNA. Hybridization was repeated three times to expand the barcoding space and ligate cell barcodes to cDNA and chromatin fragments. Reverse crosslinking was performed to release barcoded molecules. cDNA was separated from chromatin using streptavidin beads, and each library was prepared separately for sequencing.

### Quantification and sequencing

Both scATAC-seq and scRNA-seq libraries were quantified with the KAPA Library Quantification Kit and pooled for sequencing. Single cell libraries were sequenced on the Nova-seq platform (Illumina) using a 200-cycle kit (Read 1: 50 cycles, Index 1: 99 cycles, Index 2: 8 cycles, Read 2: 50 cycles). Bulk libraries were sequenced on the Nova-seq platform (Illumina) using a 100-cycle kit 483 (Read 1: 50 cycles, Index 1: 8 cycles, Index 2: 8 cycles, Read 2: 50 cycles).

### SHARE-seq data pre-processing

SHARE-seq data were processed using the SHARE-seqV2 alignment pipeline (https://github.com/masai1116/SHARE-seq-alignmentV2/) and aligned to hg38. RNA cell profiles were filtered to have at least 500 unique features (genes) and less than 5% mitochondrial reads. Seurat V3^63^ was used to scale the gene expression matrix by total UMI counts, multiplied by the mean number of transcripts, and values were log transformed. Open chromatin region peaks were called on individual samples using MACS2 peak caller^64^ with the following parameters: --nomodel – nolambda –keep-dup -call-summits. Peaks from all samples were merged and peaks overlapping with ENCODE blacklisted regions (https://sites.google.com/site/anshulkundaje/projects/blacklists) were filtered out. Peak summits were extended by 150 bp on each side and defined as accessible regions (for footprinting analyses, these peaks were later resized to 1000 bp in width). Peaks were annotated to genes using Homer^65^. The fragment counts in peaks and TF scores were calculated using chromVAR^27^. Cell barcodes with less than 45% reads in peaks (FRiP) or 2000 unique fragments in peaks were removed. The aligned reads were then intersected with peak window regions, producing a matrix of chromatin accessibility counts in peaks (rows) by cells (columns). For visualization of T cell exhaustion progression, the top 2,000 variable genes were used for principal component analysis. These principal components were batch-corrected by donor and cell cycle phase using Harmony^21^. The top 20 harmony-corrected principal components were projected into 2D space by UMAP. For all sequencing analysis after harmony-based batch correction, individual cells across donors were weighted equally and donor-blind.

### Gene modules

The top variable genes across T cell clusters in UMAP space were calculated using Seurat V3 and filtered for less than 10^-100^ adjusted P value. Expression of these genes was aggregated across the same T cell clusters to produce pseudobulks as input for DESeq2 normalization^66^. Spearman correlation was used on normalized variable gene counts to produce a distances matrix between all variable genes. Hierarchical clustering was performed on this distances matrix to create a dendrogram linking groups of co-varying genes. The number of modules produced by the tree-cutting method was determined to be 4 using the elbow plot method, minimizing the total Euclidean distance of cells in the same cluster. A combined effector – exhaustion score was created by subtracting the scaled effector module from the scaled exhaustion module. This combined score was used as a metric for exhaustion progression.

### Causal reasoning analysis

Causal reasoning analysis was performed on gene expression change associated with Iberdomide treatment as previously described^67^. The functions implementing Causal reasoning are embedded in a commercially available R toolkit called Computational Biology Methods for Drug Discovery (CBDD, Clarivate Analytics, v16.1.0). MetaBase, a manually curated database (also from Clarivate) was used as the knowledge base for drawing causal inferences. The most significant differentially expressed genes (fold change +/- 1.5, adjusted p value < 0.05) were used as input to the causal reasoning wrapper function within the toolkit. Transcription factors from the results of causal reasoning analyses were prioritized based on relevance to current biology.

### TF footprinting analysis

To comprehensively identify transcription factors (TFs) that exhibit significant changes in binding activity during a specific biological process, we employed footprinting-based analyses. We evaluated the relevance of a TF to a set of regions of interest by examining its motif enrichment and differential footprints. For the TF binding motif enrichment analysis, we utilized pycisTarget^68^ and employed the normalized enrichment score (NES) score from the software output as the quantitative metric. Notably, in the pycisTarget analysis using the SCENIC+ motif collection, each motif corresponds to a TF cluster, which can comprise one or more TFs. For the differential analysis based on footprints, we first obtained the position weight matrix (PWM) matrices for each TF cluster utilized in pycisTarget and SCENIC+. We then located the matches of a given TF cluster motif on the genome using MOODS^69^. Subsequently, we calculated the TF binding scores (TFBS) for all matches using the PRINT^45^ framework for each condition separately. We further assessed the significance of differential TFBS between the two conditions using the Wilcoxon rank-sum test. To further elucidate which TF within a SCENIC+ TF cluster might account for the enrichment of the corresponding TF cluster motif in open chromatin regions, we conducted a correlation-based analysis. Utilizing chromVAR^27^, we calculated a matrix where each entry represents the normalized enrichment of each motif in each cell. For all TFs associated with a TF cluster motif, those with a zero read count based on scRNA-seq were excluded. Subsequently, we determined the correlation between the expression level of a TF and the chromVAR score of the motif across cells. TFs with absolute correlation values equal to or greater than 90% of the maximal absolute correlation for each motif cluster were identified as the representative TFs for their respective clusters.

To demonstrate the effects of iberdomide treatment, we generated aggregate multi-scale footprints for IKZF1 binding sites. Based on MOODS^69^, we identified matches of IKZF1 motifs across genome. These matches were then categorized as either binding or non-binding sites based on the IKZF1 ChIP-seq data, employing cutoffs at the top and bottom 25% quantiles. The multi-scale footprints of treated and nontreated cells at binding and non-binding IKZF1 motif sites were aggregated into 4 maps accordingly. We normalized the IKZF1 binding footprints by subtracting the matched non-binding footprint. To discern the effect of the treatment, we calculated the differential between the normalized iberdomide treated footprint and the DMSO control.

## Acknowledgments

We thank members of AstraZeneca and the Buenrostro lab for critical reading of the manuscript and helpful discussions. We are grateful to the Broad Walkup Sequencing Core and the Bauer Core at Harvard for providing sequencing services. J.D.B. and the Buenrostro lab acknowledge support from the NIH New Innovator Award (DP2) and AstraZeneca. T.T. is supported by an NSF Fellowship.

## Author Contributions

J.K and M.G.O. generated the viral peptide cell line and developed a co-culture assay to measure tumor cell killing. A.D. and I.K.R. developed the antigen-specific exhaustion assay. G.B., I.K.R and L.P. adapted the protocol with tetramer sorting. G.B. designed and coordinated the CRISPR screen run from Astrazeneca while S.L.T. and V.B.W. ran the CRISPR screen at Revvity. E.J. and A.G. analyzed the screen dataset. E.J. ran assays testing iberdomide. I.K.R, E.J., G.B. and L.P. generated T cell samples for sequencing. T.T generated sequencing datasets. T.T. analyzed the sequencing data with computational support from R.Z., Y.H., and S.C. T.T., G.B., J.D.B., and D.A.M. wrote the manuscript. J.D.B. and D.A.M. supervised the research, and all authors reviewed the manuscript.

## References

1. Tang L, Zhang Y, Hu Y, Mei H. T Cell Exhaustion and CAR-T Immunotherapy in Hematological Malignancies. Biomed Res Int. 2021 Feb 25;2021:6616391. doi: 10.1155/2021/6616391. PMID: 33728333; PMCID: PMC7936901.

2. Gumber D, Wang LD. Improving CAR-T immunotherapy: Overcoming the challenges of T cell exhaustion. EBioMedicine. 2022 Mar;77:103941. doi: 10.1016/j.ebiom.2022.103941. Epub 2022 Mar 15. PMID: 35301179; PMCID: PMC8927848.

3. Poorebrahim M, Melief J, Pico de Coaña Y, L Wickström S, Cid-Arregui A, Kiessling R. Counteracting CAR T cell dysfunction. Oncogene. 2021 Jan;40(2):421–435. doi: 10.1038/s41388-020-01501-x. Epub 2021 Jan 14. PMID: 33168929; PMCID: PMC7808935.

4. Friedrich MJ, Neri P, Kehl N, Michel J, Steiger S, Kilian M, Leblay N, Maity R, Sankowski R, Lee H, Barakat E, Ahn S, Weinhold N, Rippe K, Bunse L, Platten M, Goldschmidt H, Müller-Tidow C, Raab MS, Bahlis NJ. The pre-existing T cell landscape determines the response to bispecific T cell engagers in multiple myeloma patients. Cancer Cell. 2023 Apr 10;41(4):711–725.e6. doi: 10.1016/j.ccell.2023.02.008. Epub 2023 Mar 9. PMID: 36898378.

5. Ghoneim HE, Fan Y, Moustaki A, Abdelsamed HA, Dash P, Dogra P, Carter R, Awad W, Neale G, Thomas PG, Youngblood B. De Novo Epigenetic Programs Inhibit PD-1 Blockade-Mediated T Cell Rejuvenation. Cell. 2017 Jun 29;170(1):142–157.e19. doi: 10.1016/j.cell.2017.06.007. Epub 2017 Jun 22. PMID: 28648661; PMCID: PMC5568784.

6. Sen DR, Kaminski J, Barnitz RA, Kurachi M, Gerdemann U, Yates KB, Tsao HW, Godec J, LaFleur MW, Brown FD, Tonnerre P, Chung RT, Tully DC, Allen TM, Frahm N, Lauer GM, Wherry EJ, Yosef N, Haining WN. The epigenetic landscape of T cell exhaustion. Science. 2016 Dec 2;354(6316):1165–1169. doi: 10.1126/science.aae0491. Epub 2016 Oct 27. PMID: 27789799; PMCID: PMC5497589.

7. Pauken KE, Sammons MA, Odorizzi PM, Manne S, Godec J, Khan O, Drake AM, Chen Z, Sen DR, Kurachi M, Barnitz RA, Bartman C, Bengsch B, Huang AC, Schenkel JM, Vahedi G, Haining WN, Berger SL, Wherry EJ. Epigenetic stability of exhausted T cells limits durability of reinvigoration by PD-1 blockade. Science. 2016 Dec 2;354(6316):1160–1165. doi: 10.1126/science.aaf2807. Epub 2016 Oct 27. PMID: 27789795; PMCID: PMC5484795.

8. Giles, Josephine R et al. “Shared and distinct biological circuits in effector, memory and exhausted CD8+ T cells revealed by temporal single-cell transcriptomics and epigenetics.” Nature immunology vol. 23,11 (2022): 1600–1613. doi:10.1038/s41590-022-01338-4

9. Miller BC, Sen DR, Al Abosy R, Bi K, Virkud YV, LaFleur MW, Yates KB, Lako A, Felt K, Naik GS, Manos M, Gjini E, Kuchroo JR, Ishizuka JJ, Collier JL, Griffin GK, Maleri S, Comstock DE, Weiss SA, Brown FD, Panda A, Zimmer MD, Manguso RT, Hodi FS, Rodig SJ, Sharpe AH, Haining WN. Subsets of exhausted CD8+ T cells differentially mediate tumor control and respond to checkpoint blockade. Nat Immunol. 2019 Mar;20(3):326–336. doi: 10.1038/s41590-019-0312-6. Epub 2019 Feb 18. Erratum in: Nat Immunol. 2019 Nov;20(11):1556. PMID: 30778252; PMCID: PMC6673650.

10. Zhang F, Zhou X, DiSpirito JR, Wang C, Wang Y, Shen H. Epigenetic manipulation restores functions of defective CD8⁺ T cells from chronic viral infection. Mol Ther. 2014 Sep;22(9):1698–706. doi: 10.1038/mt.2014.91. Epub 2014 May 27. PMID: 24861055; PMCID: PMC4435497.

11. Laino AS, Betts BC, Veerapathran A, Dolgalev I, Sarnaik A, Quayle SN, Jones SS, Weber JS, Woods DM. HDAC6 selective inhibition of melanoma patient T-cells augments anti-tumor characteristics. J Immunother Cancer. 2019 Feb 6;7(1):33. doi: 10.1186/s40425-019-0517-0. PMID: 30728070; PMCID: PMC6366050.

12. Battistello E, Hixon KA, Comstock DE, Collings CK, Chen X, Rodriguez Hernaez J, Lee S, Cervantes KS, Hinkley MM, Ntatsoulis K, Cesarano A, Hockemeyer K, Haining WN, Witkowski MT, Qi J, Tsirigos A, Perna F, Aifantis I, Kadoch C. Stepwise activities of mSWI/SNF family chromatin remodeling complexes direct T cell activation and exhaustion. Mol Cell. 2023 Apr 20;83(8):1216–1236.e12. doi: 10.1016/j.molcel.2023.02.026. Epub 2023 Mar 20. PMID: 36944333; PMCID: PMC10121856.

13. Belk JA, Yao W, Ly N, Freitas KA, Chen YT, Shi Q, Valencia AM, Shifrut E, Kale N, Yost KE, Duffy CV, Daniel B, Hwee MA, Miao Z, Ashworth A, Mackall CL, Marson A, Carnevale J, Vardhana SA, Satpathy AT. Genome-wide CRISPR screens of T cell exhaustion identify chromatin remodeling factors that limit T cell persistence. Cancer Cell. 2022 Jul 11;40(7):768–786.e7. doi: 10.1016/j.ccell.2022.06.001. Epub 2022 Jun 23. PMID: 35750052; PMCID: PMC9949532.

14. Blake MK, O’Connell P, Aldhamen YA. Fundamentals to therapeutics: Epigenetic modulation of CD8+ T Cell exhaustion in the tumor microenvironment. Front Cell Dev Biol. 2023 Jan 4;10:1082195. doi: 10.3389/fcell.2022.1082195. PMID: 36684449; PMCID: PMC9846628.

15. Balkhi MY, Wittmann G, Xiong F, Junghans RP. YY1 Upregulates Checkpoint Receptors and Downregulates Type I Cytokines in Exhausted, Chronically Stimulated Human T Cells. iScience. 2018 Apr 27;2:105–122. doi: 10.1016/j.isci.2018.03.009. Epub 2018 Apr 11. PMID: 30428369; PMCID: PMC6136936.

16. Hou Y, Zak J, Shi Y, Pratumchai I, Dinner B, Wang W, Qin K, Weber E, Teijaro JR, Wu P. Transient EZH2 suppression by Tazemetostat during in vitro expansion maintains T cell stemness and improves adoptive T cell therapy. bioRxiv [Preprint]. 2023 Feb 7:2023.02.07.527459. doi: 10.1101/2023.02.07.527459. PMID: 36798389; PMCID: PMC9934551.

17. Lonial S, Popat R, Hulin C, Jagannath S, Oriol A, Richardson PG, Facon T, Weisel K, Larsen JT, Minnema MC, Abdallah AO, Badros AZ, Knop S, Stadtmauer EA, Cheng Y, Amatangelo M, Chen M, Nguyen TV, Amin A, Peluso T, van de Donk NWCJ. Iberdomide plus dexamethasone in heavily pretreated late-line relapsed or refractory multiple myeloma (CC-220-MM-001): a multicentre, multicohort, open-label, phase 1/2 trial. Lancet Haematol. 2022 Nov;9(11):e822–e832. doi: 10.1016/S2352-3026(22)00290-3. Epub 2022 Oct 6. PMID: 36209764.

18. Saadey AA, Yousif A, Osborne N, Shahinfar R, Chen YL, Laster B, Rajeev M, Bauman P, Webb A, Ghoneim HE. Rebalancing TGFβ1/BMP signals in exhausted T cells unlocks responsiveness to immune checkpoint blockade therapy. Nat Immunol. 2023 Feb;24(2):280–294. doi: 10.1038/s41590-022-01384-y. Epub 2022 Dec 21. PMID: 36543960.

19. Dunsford, L.S., Thoirs, R.H., Rathbone, E., and Patakas, A. (2020). A human in vitro T cell exhaustion model for assessing immuno-oncology therapies. pp. 89–101. doi: 10.1007/978-1-0716-0171-6_6.

20. Ma S, Zhang B, LaFave LM, Earl AS, Chiang Z, Hu Y, Ding J, Brack A, Kartha VK, Tay T, Law T, Lareau C, Hsu YC, Regev A, Buenrostro JD. Chromatin Potential Identified by Shared Single-Cell Profiling of RNA and Chromatin. Cell. 2020 Nov 12;183(4):1103–1116.e20. doi: 10.1016/j.cell.2020.09.056. Epub 2020 Oct 23. PMID: 33098772; PMCID: PMC7669735.

21. Korsunsky I, Millard N, Fan J, Slowikowski K, Zhang F, Wei K, Baglaenko Y, Brenner M, Loh PR, Raychaudhuri S. Fast, sensitive and accurate integration of single-cell data with Harmony. Nat Methods. 2019 Dec;16(12):1289–1296. doi: 10.1038/s41592-019-0619-0. Epub 2019 Nov 18. PMID: 31740819; PMCID: PMC6884693.

22. Kumar J, Kumar R, Kumar Singh A, Tsakem EL, Kathania M, Riese MJ, Theiss AL, Davila ML, Venuprasad K. Deletion of Cbl-b inhibits CD8+ T-cell exhaustion and promotes CAR T-cell function. J Immunother Cancer. 2021 Jan;9(1):e001688. doi: 10.1136/jitc-2020-001688. PMID: 33462140; PMCID: PMC7813298.

23. Trefny MP, Kirchhammer N, Auf der Maur P, Natoli M, Schmid D, Germann M, Fernandez Rodriguez L, Herzig P, Lötscher J, Akrami M, Stinchcombe JC, Stanczak MA, Zingg A, Buchi M, Roux J, Marone R, Don L, Lardinois D, Wiese M, Jeker LT, Bentires-Alj M, Rossy J, Thommen DS, Griffiths GM, Läubli H, Hess C, Zippelius A. Deletion of SNX9 alleviates CD8 T cell exhaustion for effective cellular cancer immunotherapy. Nat Commun. 2023 Feb 2;14(1):86. doi: 10.1038/s41467-022-35583-w. PMID: 36732507; PMCID: PMC9895440.

24. Gupta PK, Godec J, Wolski D, Adland E, Yates K, Pauken KE, Cosgrove C, Ledderose C, Junger WG, Robson SC, Wherry EJ, Alter G, Goulder PJ, Klenerman P, Sharpe AH, Lauer GM, Haining WN. CD39 Expression Identifies Terminally Exhausted CD8+ T Cells. PLoS Pathog. 2015 Oct 20;11(10):e1005177. doi: 10.1371/journal.ppat.1005177. PMID: 26485519; PMCID: PMC4618999.

25. Zheng L, Qin S, Si W, Wang A, Xing B, Gao R, Ren X, Wang L, Wu X, Zhang J, Wu N, Zhang N, Zheng H, Ouyang H, Chen K, Bu Z, Hu X, Ji J, Zhang Z. Pan-cancer single-cell landscape of tumor-infiltrating T cells. Science. 2021 Dec 17;374(6574):abe6474. doi: 10.1126/science.abe6474. Epub 2021 Dec 17. PMID: 34914499

26. Ou R, Zhang M, Huang L, Moskophidis D. Control of virus-specific CD8+ T-cell exhaustion and immune-mediated pathology by E3 ubiquitin ligase Cbl-b during chronic viral infection. J Virol. 2008 Apr;82(7):3353–68. doi: 10.1128/JVI.01350-07. Epub 2008 Jan 16. PMID: 18199651; PMCID: PMC2268476.

27. Schep AN, Wu B, Buenrostro JD, Greenleaf WJ. chromVAR: inferring transcription-factor-associated accessibility from single-cell epigenomic data. Nat Methods. 2017 Oct;14(10):975–978. doi: 10.1038/nmeth.4401. Epub 2017 Aug 21. PMID: 28825706; PMCID: PMC5623146.

28. Nah J, Seong RH. Krüppel-like factor 4 regulates the cytolytic effector function of exhausted CD8 T cells. Sci Adv. 2022 Nov 25;8(47):eadc9346. doi: 10.1126/sciadv.adc9346. Epub 2022 Nov 25. PMID: 36427304; PMCID: PMC9699681.

29. Seo W, Jerin C, Nishikawa H. Transcriptional regulatory network for the establishment of CD8+ T cell exhaustion. Exp Mol Med. 2021 Feb;53(2):202–209. doi: 10.1038/s12276-021-00568-0. Epub 2021 Feb 24. PMID: 33627794; PMCID: PMC8080584.

30. Delpoux A, Marcel N, Hess Michelini R, Katayama CD, Allison KA, Glass CK, Quiñones-Parra SM, Murre C, Loh L, Kedzierska K, Lappas M, Hedrick SM, Doedens AL. FOXO1 constrains activation and regulates senescence in CD8 T cells. Cell Rep. 2021 Jan 26;34(4):108674. doi: 10.1016/j.celrep.2020.108674. PMID: 33503413.

31. O’Brien S, Thomas RM, Wertheim GB, Zhang F, Shen H, Wells AD. IKAROS imposes a barrier to CD8+ T cell differentiation by restricting autocrine IL-2 production. J Immunol. 2014 Jun 1;192(11):5118–29. doi: 10.4049/jimmunol.1301992. Epub 2014 Apr 28. PMID: 24778448; PMCID: PMC4042307.

32. Prinzing B, Zebley CC, Petersen CT, Fan Y, Anido AA, Yi Z, Nguyen P, Houke H, Bell M, Haydar D, Brown C, Boi SK, Alli S, Crawford JC, Riberdy JM, Park JJ, Zhou S, Velasquez MP, DeRenzo C, Lazzarotto CR, Tsai SQ, Vogel P, Pruett-Miller SM, Langfitt DM, Gottschalk S, Youngblood B, Krenciute G. Deleting DNMT3A in CAR T cells prevents exhaustion and enhances antitumor activity. Sci Transl Med. 2021 Nov 17;13(620):eabh0272. doi: 10.1126/scitranslmed.abh0272. Epub 2021 Nov 17. PMID: 34788079; PMCID: PMC8733956.

33. Friend SF, Peterson LK, Treacy E, Stefanski AL, Sosinowski T, Pennock ND, Berger AJ, Winn VD, Dragone LL. The discovery of a reciprocal relationship between tyrosine-kinase signaling and cullin neddylation. PLoS One. 2013 Oct 4;8(10):e75200. doi: 10.1371/journal.pone.0075200. PMID: 24124476; PMCID: PMC3790728.

34. Friend SF, Deason-Towne F, Peterson LK, Berger AJ, Dragone LL. Regulation of T cell receptor complex-mediated signaling by ubiquitin and ubiquitin-like modifications. Am J Clin Exp Immunol. 2014 Dec 5;3(3):107–23. PMID: 25628960; PMCID: PMC4299764.

35. Lawrence T, Bebien M, Liu GY, Nizet V, Karin M. IKKalpha limits macrophage NF-kappaB activation and contributes to the resolution of inflammation. Nature. 2005 Apr 28;434(7037):1138-43. doi: 10.1038/nature03491. PMID: 15858576.

36. Liu B, Yang Y, Chernishof V, Loo RR, Jang H, Tahk S, Yang R, Mink S, Shultz D, Bellone CJ, Loo JA, Shuai K. Proinflammatory stimuli induce IKKalpha-mediated phosphorylation of PIAS1 to restrict inflammation and immunity. Cell. 2007 Jun 1;129(5):903–14. doi: 10.1016/j.cell.2007.03.056. PMID: 17540171.

37. Avitahl N, Winandy S, Friedrich C, Jones B, Ge Y, Georgopoulos K. Ikaros sets thresholds for T cell activation and regulates chromosome propagation. Immunity. 1999 Mar;10(3):333–43. doi: 10.1016/s1074-7613(00)80033-3. PMID: 10204489.

38. Winandy S, Wu P, Georgopoulos K. A dominant mutation in the Ikaros gene leads to rapid development of leukemia and lymphoma. Cell. 1995 Oct 20;83(2):289–99. doi: 10.1016/0092-8674(95)90170-1. PMID: 7585946.

39. Surka C, Jin L, Mbong N, Lu CC, Jang IS, Rychak E, Mendy D, Clayton T, Tindall E, Hsu C, Fontanillo C, Tran E, Contreras A, Ng SWK, Matyskiela M, Wang K, Chamberlain P, Cathers B, Carmichael J, Hansen J, Wang JCY, Minden MD, Fan J, Pierce DW, Pourdehnad M, Rolfe M, Lopez-Girona A, Dick JE, Lu G. CC-90009, a novel cereblon E3 ligase modulator, targets acute myeloid leukemia blasts and leukemia stem cells. Blood. 2021 Feb 4;137(5):661–677. doi: 10.1182/blood.2020008676. PMID: 33197925; PMCID: PMC8215192.

40. Watson ER, Novick S, Matyskiela ME, Chamberlain PP, H de la Peña A, Zhu J, Tran E, Griffin PR, Wertz IE, Lander GC. Molecular glue CELMoD compounds are regulators of cereblon conformation. Science. 2022 Nov 4;378(6619):549–553. doi: 10.1126/science.add7574. Epub 2022 Nov 3. PMID: 36378961; PMCID: PMC9714526.

41. Ping N, Qu C, Li M, Kang L, Kong D, Chen X, Wu Q, Xia F, Yu L, Yao H, Yan L, Wu D, Jin Z. Overall survival benefits provided by lenalidomide maintenance after chimeric antigen receptor T cell therapy in patients with refractory/relapsed diffuse large B-cell lymphoma. Ann Transl Med. 2022 Mar;10(6):298. doi: 10.21037/atm-22-20. PMID: 35433994; PMCID: PMC9011311.

42. Matyskiela ME, Zhang W, Man HW, Muller G, Khambatta G, Baculi F, Hickman M, LeBrun L, Pagarigan B, Carmel G, Lu CC, Lu G, Riley M, Satoh Y, Schafer P, Daniel TO, Carmichael J, Cathers BE, Chamberlain PP. A Cereblon Modulator (CC-220) with Improved Degradation of IKAROS and Aiolos. J Med Chem. 2018 Jan 25;61(2):535–542. doi: 10.1021/acs.jmedchem.6b01921. Epub 2017 Apr 20. PMID: 28425720.

43. Chen J, López-Moyado IF, Seo H, Lio CJ, Hempleman LJ, Sekiya T, Yoshimura A, Scott-Browne JP, Rao A. NR4A transcription factors limit CAR T cell function in solid tumours. Nature. 2019 Mar;567(7749):530–534. doi: 10.1038/s41586-019-0985-x. Epub 2019 Feb 27. PMID: 30814732; PMCID: PMC6546093.

44. Chindelevitch L, Ziemek D, Enayetallah A, Randhawa R, Sidders B, Brockel C, Huang ES. Causal reasoning on biological networks: interpreting transcriptional changes. Bioinformatics. 2012 Apr 15;28(8):1114–21. doi: 10.1093/bioinformatics/bts090. Epub 2012 Feb 21. PMID: 22355083.

45. Zhong Y, Walker SK, Pritykin Y, Leslie CS, Rudensky AY, van der Veeken J. Hierarchical regulation of the resting and activated T cell epigenome by major transcription factor families. Nat Immunol. 2022 Jan;23(1):122–134. doi: 10.1038/s41590-021-01086-x. Epub 2021 Dec 22. PMID: 34937932; PMCID: PMC8712421.

46. Buenrostro JD, Giresi PG, Zaba LC, Chang HY, Greenleaf WJ. Transposition of native chromatin for fast and sensitive epigenomic profiling of open chromatin, DNA-binding proteins and nucleosome position. Nat Methods. 2013 Dec;10(12):1213–8. doi: 10.1038/nmeth.2688. Epub 2013 Oct 6. PMID: 24097267; PMCID: PMC3959825.

47. Zheng, Rongbin, et al. “Cistrome Data Browser: expanded datasets and new tools for gene regulatory analysis.” Nucleic acids research vol. 47,D1 (2019): D729–D735. doi:10.1093/nar/gky1094

48. Hu Y, Ma S, Kartha VK, Duarte FM, Horlbeck M, Zhang R, Shrestha R, Labade A, Kletzien H, Meliki A, Castillo A, Durand N, Mattei E, Anderson LJ, Tay T, Earl AS, Shoresh N, Epstein CB, Wagers A, Buenrostro JD. Single-cell multi-scale footprinting reveals the modular organization of DNA regulatory elements. bioRxiv [Preprint]. 2023 Mar 29:2023.03.28.533945. doi: 10.1101/2023.03.28.533945. PMID: 37034577; PMCID: PMC10081223.

49. Köpf-Maier P, Sass G. Antitumor activity of treosulfan against human breast carcinomas. Cancer Chemother Pharmacol. 1992;31(2):103–10. doi: 10.1007/BF00685095. PMID: 1451231.

50. Perotti EA, Georgopoulos K, Yoshida T. An Ikaros Promoter Element with Dual Epigenetic and Transcriptional Activities. PLoS One. 2015 Jul 2;10(7):e0131568. doi: 10.1371/journal.pone.0131568. PMID: 26135129; PMCID: PMC4489883.

51. Trinh LA, Ferrini R, Cobb BS, Weinmann AS, Hahm K, Ernst P, Garraway IP, Merkenschlager M, Smale ST. Down-regulation of TDT transcription in CD4(+)CD8(+) thymocytes by Ikaros proteins in direct competition with an Ets activator. Genes Dev. 2001 Jul 15;15(14):1817–32. doi: 10.1101/gad.905601. PMID: 11459831; PMCID: PMC312741.

52. Sun W, Guo J, McClellan D, Poeschla A, Bareyan D, Casey MJ, Cairns BR, Tantin D, Engel ME. GFI1 Cooperates with IKZF1/IKAROS to Activate Gene Expression in T-cell Acute Lymphoblastic Leukemia. Mol Cancer Res. 2022 Apr 1;20(4):501–514. doi: 10.1158/1541-7786.MCR-21-0352. PMID: 34980595; PMCID: PMC8983472.

53. Oravecz, Attila et al. “Ikaros mediates gene silencing in T cells through Polycomb repressive complex 2.” Nature communications vol. 6 8823. 9 Nov. 2015, doi:10.1038/ncomms9823

54. Powell, Michael D et al. “Ikaros Zinc Finger Transcription Factors: Regulators of Cytokine Signaling Pathways and CD4+ T Helper Cell Differentiation.” Frontiers in immunology vol. 10 1299. 6 Jun. 2019, doi:10.3389/fimmu.2019.01299

55. Lyon de Ana, Carolina, et al. “Lack of Ikaros Deregulates Inflammatory Gene Programs in T Cells.” Journal of immunology (Baltimore, Md. : 1950) vol. 202,4 (2019): 1112–1123. doi:10.4049/jimmunol.1801270

56. Koipally, J et al. “Repression by Ikaros and Aiolos is mediated through histone deacetylase complexes.” The EMBO journal vol. 18,11 (1999): 3090–100. doi:10.1093/emboj/18.11.3090

57. Bandyopadhyay, Sanmay et al. “Interleukin 2 gene transcription is regulated by Ikaros-induced changes in histone acetylation in anergic T cells.” Blood vol. 109,7 (2007): 2878–86. doi:10.1182/blood-2006-07-037754

58. Hay, J, et al. “Abstract 5586: Demonstrating restoration of T cell function in exhausted T cells with IKZF3 small molecule inhibitor, Lenalidomide.” Cancer Res vol. 82,12_Supplement (2022): 5586. doi:10.1158/1538-7445.AM2022-5586

59. Zou, Yan et al. “IKZF3 deficiency potentiates chimeric antigen receptor T cells targeting solid tumors.” Cancer letters vol. 524 (2022): 121–130. doi:10.1016/j.canlet.2021.10.016

60. Ye, Ying et al. “First-in-Human, Single- and Multiple-Ascending-Dose Studies in Healthy Subjects to Assess Pharmacokinetics, Pharmacodynamics, and Safety/Tolerability of Iberdomide, a Novel Cereblon E3 Ligase Modulator.” Clinical pharmacology in drug development vol. 10,5 (2021): 471–485. doi:10.1002/cpdd.869

61. D’Souza, Criselle et al. “Understanding the Role of T-Cells in the Antimyeloma Effect of Immunomodulatory Drugs.” Frontiers in immunology vol. 12 632399. 5 Mar. 2021, doi:10.3389/fimmu.2021.632399

62. Correa, Luis O et al. “DNA Methylation in T-Cell Development and Differentiation.” Critical reviews in immunology vol. 40,2 (2020): 135–156. doi:10.1615/CritRevImmunol.2020033728

63. Zhang, Qing et al. “Histone Deacetylases (HDACs) Guided Novel Therapies for T-cell lymphomas.” International journal of medical sciences vol. 16,3 424–442. 29 Jan. 2019, doi:10.7150/ijms.30154

64. Zhong, Mengjun et al. “BET bromodomain inhibition rescues PD-1-mediated T-cell exhaustion in acute myeloid leukemia.” Cell death & disease vol. 13,8 671. 2 Aug. 2022, doi:10.1038/s41419-022-05123-x

65. Kong, Weimin et al. “BET bromodomain protein inhibition reverses chimeric antigen receptor extinction and reinvigorates exhausted T cells in chronic lymphocytic leukemia.” The Journal of clinical investigation vol. 131,16 (2021): e145459. doi:10.1172/JCI145459

66. Seo H, Chen J, González-Avalos E, Samaniego-Castruita D, Das A, Wang YH, López-Moyado IF, Georges RO, Zhang W, Onodera A, Wu CJ, Lu LF, Hogan PG, Bhandoola A, Rao A. TOX and TOX2 transcription factors cooperate with NR4A transcription factors to impose CD8+ T cell exhaustion. Proc Natl Acad Sci U S A. 2019 Jun 18;116(25):12410–12415. doi: 10.1073/pnas.1905675116. Epub 2019 May 31. Erratum in: Proc Natl Acad Sci U S A. 2019 Sep 24;116(39):19761. PMID: 31152140; PMCID: PMC6589758.

67. Khan O, Giles JR, McDonald S, Manne S, Ngiow SF, Patel KP, Werner MT, Huang AC, Alexander KA, Wu JE, Attanasio J, Yan P, George SM, Bengsch B, Staupe RP, Donahue G, Xu W, Amaravadi RK, Xu X, Karakousis GC, Mitchell TC, Schuchter LM, Kaye J, Berger SL, Wherry EJ. TOX transcriptionally and epigenetically programs CD8+ T cell exhaustion. Nature. 2019 Jul;571(7764):211–218. doi: 10.1038/s41586-019-1325-x. Epub 2019 Jun 17. PMID: 31207603; PMCID: PMC6713202.

68. Scott AC, Dündar F, Zumbo P, Chandran SS, Klebanoff CA, Shakiba M, Trivedi P, Menocal L, Appleby H, Camara S, Zamarin D, Walther T, Snyder A, Femia MR, Comen EA, Wen HY, Hellmann MD, Anandasabapathy N, Liu Y, Altorki NK, Lauer P, Levy O, Glickman MS, Kaye J, Betel D, Philip M, Schietinger A. TOX is a critical regulator of tumour-specific T cell differentiation. Nature. 2019 Jul;571(7764):270–274. doi: 10.1038/s41586-019-1324-y. Epub 2019 Jun 17. PMID: 31207604; PMCID: PMC7698992.

69. Yao C, Sun HW, Lacey NE, Ji Y, Moseman EA, Shih HY, Heuston EF, Kirby M, Anderson S, Cheng J, Khan O, Handon R, Reilley J, Fioravanti J, Hu J, Gossa S, Wherry EJ, Gattinoni L, McGavern DB, O’Shea JJ, Schwartzberg PL, Wu T. Single-cell RNA-seq reveals TOX as a key regulator of CD8+ T cell persistence in chronic infection. Nat Immunol. 2019 Jul;20(7):890–901. doi: 10.1038/s41590-019-0403-4. Epub 2019 Jun 17. PMID: 31209400; PMCID: PMC6588409.

70. Stuart, Tim et al. “Comprehensive Integration of Single-Cell Data.” Cell vol. 177,7 (2019): 1888–1902.e21. doi:10.1016/j.cell.2019.05.031

71. Zhang, Yong et al. “Model-based analysis of ChIP-Seq (MACS).” Genome biology vol. 9,9 (2008): R137. doi:10.1186/gb-2008-9-9-r137

72. Heinz, Sven et al. “Simple combinations of lineage-determining transcription factors prime cis-regulatory elements required for macrophage and B cell identities.” Molecular cell vol. 38,4 (2010): 576–89. doi:10.1016/j.molcel.2010.05.004

73. Love, Michael I et al. “Moderated estimation of fold change and dispersion for RNA-seq data with DESeq2.” Genome biology vol. 15,12 (2014): 550. doi:10.1186/s13059-014-0550-8

74. Criscione, Steven W et al. “The landscape of therapeutic vulnerabilities in EGFR inhibitor osimertinib drug tolerant persister cells.” NPJ precision oncology vol. 6,1 95. 27 Dec. 2022, doi:10.1038/s41698-022-00337-w

75. Bravo González-Blas, Carmen et al. “SCENIC+: single-cell multiomic inference of enhancers and gene regulatory networks.” Nature methods vol. 20,9 (2023): 1355–1367. doi:10.1038/s41592-023-01938-4

76. Korhonen, Janne et al. “MOODS: fast search for position weight matrix matches in DNA sequences.” Bioinformatics (Oxford, England) vol. 25,23 (2009): 3181–2. doi:10.1093/bioinformatics/btp554

77. Sumida TS, Dulberg S, Schupp JC, Lincoln MR, Stillwell HA, Axisa PP, Comi M, Unterman A, Kaminski N, Madi A, Kuchroo VK, Hafler DA. Type I interferon transcriptional network regulates expression of coinhibitory receptors in human T cells. Nat Immunol. 2022 Apr;23(4):632–642. doi: 10.1038/s41590-022-01152-y. Epub 2022 Mar 17. PMID: 35301508; PMCID: PMC8989655.

78. Bachireddy, Pavan et al. “Mapping the evolution of T cell states during response and resistance to adoptive cellular therapy.” Cell reports vol. 37,6 (2021): 109992. doi:10.1016/j.celrep.2021.109992

79. Alfei F, Kanev K, Hofmann M, Wu M, Ghoneim HE, Roelli P, Utzschneider DT, von Hoesslin M, Cullen JG, Fan Y, Eisenberg V, Wohlleber D, Steiger K, Merkler D, Delorenzi M, Knolle PA, Cohen CJ, Thimme R, Youngblood B, Zehn D. TOX reinforces the phenotype and longevity of exhausted T cells in chronic viral infection. Nature. 2019 Jul;571(7764):265–269. doi: 10.1038/s41586-019-1326-9. Epub 2019 Jun 17. PMID: 31207605.

80. Liu X, Wang Y, Lu H, Li J, Yan X, Xiao M, Hao J, Alekseev A, Khong H, Chen T, Huang R, Wu J, Zhao Q, Wu Q, Xu S, Wang X, Jin W, Yu S, Wang Y, Wei L, Wang A, Zhong B, Ni L, Liu X, Nurieva R, Ye L, Tian Q, Bian XW, Dong C. Genome-wide analysis identifies NR4A1 as a key mediator of T cell dysfunction. Nature. 2019 Mar;567(7749):525–529. doi: 10.1038/s41586-019-0979-8. Epub 2019 Feb 27. PMID: 30814730; PMCID: PMC6507425.

81. Wang JH, Avitahl N, Cariappa A, Friedrich C, Ikeda T, Renold A, Andrikopoulos K, Liang L, Pillai S, Morgan BA, Georgopoulos K. Aiolos regulates B cell activation and maturation to effector state. Immunity. 1998 Oct;9(4):543–53. doi: 10.1016/s1074-7613(00)80637-8. PMID: 9806640.

